# Histone acetylome-wide association study of tuberculosis

**DOI:** 10.1101/644112

**Authors:** Ricardo C.H. del Rosario, Jeremie Poschmann, Pavanish Kumar, Catherine Y. Cheng, Seow Theng Ong, Hajira Shreen Hajan, Dilip Kumar, Mardiana Marzuki, Xiaohua Lu, Andrea Lee, Yanxia Tang, Cynthia Bin Eng Chee, Carey Lim, Bernett Lee, Josephine Lum, Francesca Zolezzi, Michael Poidinger, Olaf Rotzschke, Chiea Chuen Khor, Yee T. Wang, K. George Chandy, Gennaro De Libero, Amit Singhal, Shyam Prabhakar

**Affiliations:** Genome Institute of Singapore, Agency for Science, Technology and Research (A*STAR), Singapore, 138672; Singapore Immunology Network, A*STAR, Singapore 138648; Lee Kong Chian School of Medicine, Nanyang Technological University Singapore, Singapore 636921; Tuberculosis control unit, Tan Tock Seng Hospital, Singapore 308087; Department of Biomedicine, University of Basel, 4056 Basel, Switzerland; Stanley Center for Psychiatric Research, Broad Institute of MIT and Harvard, 75 Ames St., Cambridge MA 02144; Centre de Recherche en Transplantation et Immunologie, UMR1064, Inserm, CHU-Nantes Université de Nantes, Nantes, France

## Abstract

Host-cell chromatin changes are thought to play an important role in the pathogenesis of infectious diseases. Here, we describe the first histone acetylome-wide association study (HAWAS) of an infectious disease, based on genome-wide H3K27 acetylation profiling of peripheral granulocytes and monocytes from subjects with active *Mycobacterium tuberculosis* (*Mtb*) infection and healthy controls. We detected >2,000 differentially acetylated loci in either cell type in a Chinese discovery cohort, which were validated in a subsequent multi-ethnic cohort, thus demonstrating that HAWAS can be independently corroborated. Acetylation changes were correlated with differential gene expression in a third cohort. Differential acetylation was enriched near potassium channel genes, including *KCNJ15*, which modulated Akt-mTOR signaling and promoted *Mtb* clearance *in vitro.* We performed histone acetylation QTL analysis on the dataset and identified candidate causal variants for immune phenotypes. Our study serves as proof-of-principle for HAWAS to infer mechanisms of host response to pathogens.

## Introduction

Tuberculosis (TB), caused by the bacterium *Mycobacterium tuberculosis* (*Mtb*), is the world’s deadliest infectious disease, causing 1.5 million deaths per year^1^. *Mtb* circumvents the immune system by remodeling the host transcriptome, leading to inhibition of protective and induction of pathological immune responses^2–4^. In particular, gene expression studies on whole blood from active TB (ATB) patients suggest that alterations in pathways such as interferon signaling, inflammation, apoptosis and pattern recognition receptor signaling may contribute to TB pathogenesis^4–6^. It is likely that these transcriptomic changes in host immune cells during infection are chromatin-mediated^7–10^.

Histone H3 acetylation at lysine 27 (H3K27ac), is a well-established chromatin signature of active enhancers and promoters^11,12^ that correlates with gene expression and transcription factor binding^13^. We previously profiled H3K27ac genome-wide in post-mortem autism spectrum disorder (ASD) brain vs. control and detected widespread ASD-associated chromatin perturbations converging upon specific pathways, thus providing a resource for subsequent mechanistic studies of ASD pathology^14^. Two other recent studies used the same methodology to detect thousands of histone acetylation changes in post-mortem brain samples from individuals with Alzheimer’s disease^15,16^. We hypothesize that the histone acetylome-wide association study (HAWAS) approach could also provide molecular insights into non-neurological conditions such as *Mtb* infection.

Here, we report a HAWAS of peripheral monocytes and granulocytes from ATB patients and age-matched healthy controls. Uniquely, we profiled histone acetylation in two distinct cohorts: discovery and validation. We also generated transcriptome data from a third cohort for further corroboration of infection-associated chromatin changes. We then analyzed the data to infer specific host-response mechanisms. Finally, we used the ChIP-seq data to call SNPs within monocyte and granulocyte regulatory elements and identify histone acetylation quantitative trait loci (haQTLs), some of which may contribute to inflammatory and infectious disease susceptibility.

## Results

### HAWAS of active Tuberculosis

We used ChIP-seq to profile H3K27ac in a total of 190 peripheral blood granulocytes and monocyte samples from ATB patients and age-matched healthy controls (HC; Fig. 1a), of which 135 were retained after QC (Fig. 1b and Supplementary Table 1). These 135 samples were split into discovery and validation cohorts, and regulatory elements in each cohort were detected as focal peaks in the ChIP-seq signal (Methods). Our granulocyte and monocyte H3K27ac profiles were highly consistent with those from the International Human Epigenome Consortium^17^ (IHEC; Extended Data Fig. 1a).

**Fig. 1.**
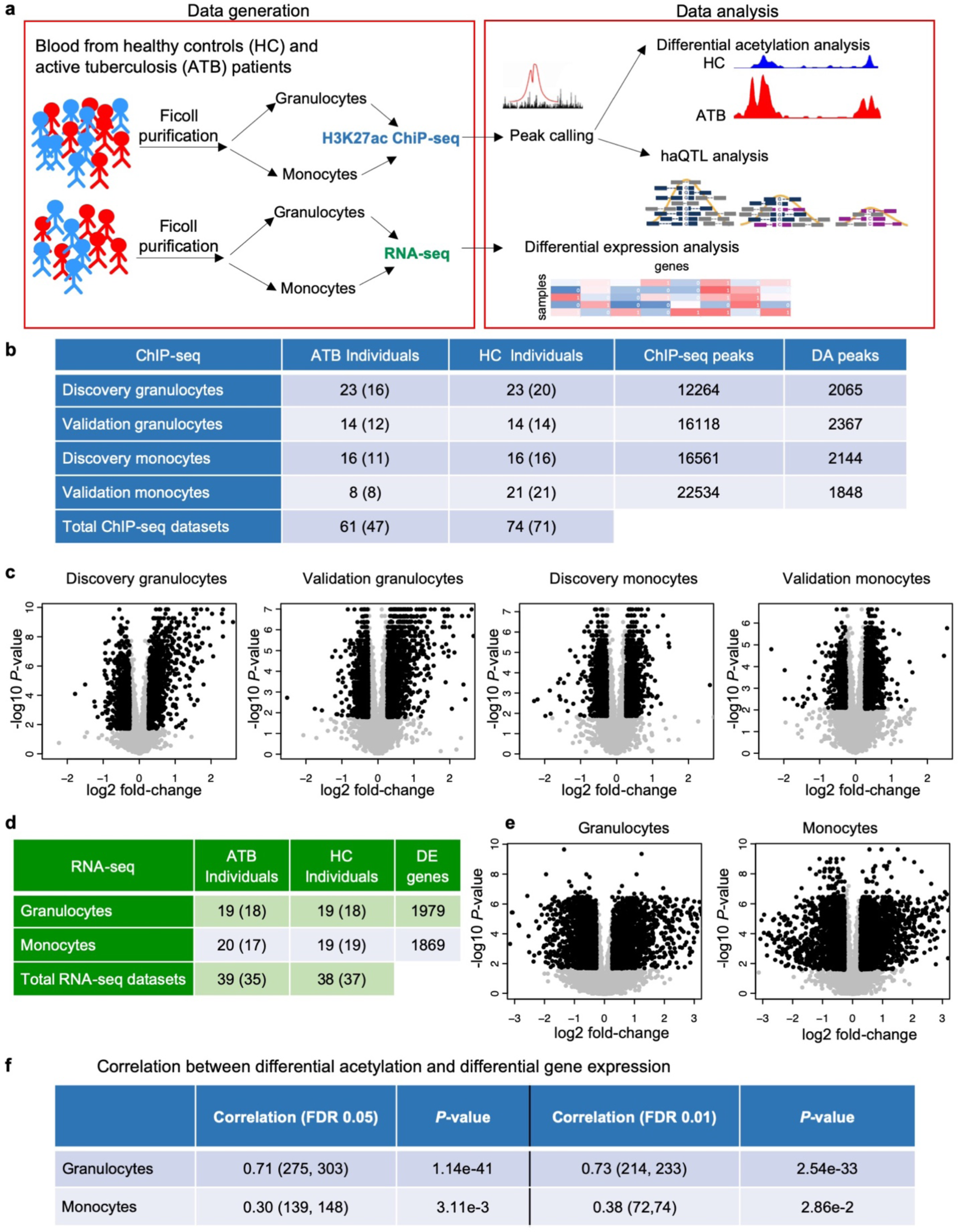
Histone acetylome-wide association study of TB. (**a**) Overview of data generation and analysis. (**b**) Number of high-quality granulocyte and monocyte ChIP-seq datasets from discovery and validation cohorts remaining after QC. Sizes of corresponding final core sets are indicated in parentheses. Number of consensus ChIP-seq peaks in high-quality datasets. Number of differential acetylated (DA) peaks in core set. (**c**) Volcano plots showing log2 fold-change and *P*-value for consensus ChIP-seq peaks (black dots: DA peaks). (**d**) Statistics of RNA-seq dataset, similar to (b). (**e**) Volcano plots showing log2 fold-change and *P*-value for expressed genes (black dots: DE genes). (**f**) Pearson correlation of log-fold-change between DE genes and their associated DA peaks. Correlations were calculated for DE and DA sets defined using the default FDR threshold (FDR≤0.05), and also a more stringent threshold (FDR≤0.01). The number of DE genes with at least one associated with DA peak and the total number of points used in the correlation are shown in parentheses. *P*-values indicate concordance of the fold-change direction, calculated using the hypergeometric test.

To identify infection-associated changes in the regulatory elements, we tested each of them for differential histone acetylation (differential peak height) between ATB and HC samples in each cohort, after controlling for confounders (Extended Data Fig. 1b-d; Methods). In total, we identified >1,800 DA peaks in each combination of cell type and cohort (Fig. 1b,c and Supplementary Table 1), indicating that ATB-associated chromatin changes were widespread in both granulocytes and monocytes. Moreover, the observed histone acetylation changes were shared across ATB individuals, rather than limited to a subset (Extended Data Fig. 2a).

Next, we investigated the correspondence between histone acetylation changes and differential gene expression. Note that, although absolute histone acetylation and gene expression readouts are strongly correlated across a single genome^13^, we expect measures of differential acetylation and differential expression to show only moderate correlation, largely due to lower signal-to-noise ratio^14^. We performed RNA-seq on peripheral granulocytes and monocytes from an independent multi-ethnic ATB vs. HC cohort and identified 1,800 differentially expressed (DE) genes in either cell type (Fig. 1d,e, Extended Data Fig. 2b and Supplementary Table 1; Methods). As expected, DE gene fold-changes in granulocytes and monocytes were significantly correlated with fold-changes of flanking DA peaks (Fig. 1f), and thus the two data types corroborated each other. However, the correlation was only moderate (granulocytes: *R*=0.71; monocytes: *R*=0.30), as was also observed in our previous HAWAS study of ASD^14^ (*R*=0.33-0.38;). These results further support our previous finding that, while differential H3K27ac ChIP-seq and transcriptome profiles tend to correlate globally, they are not mutually redundant. Rather, they provide complementary views of the disease-associated changes under investigation.

One issue not addressed in previous HAWAS studies^14–16^ is reproducibility of DA signals. First, we inspected a cluster of DA peaks detected in both granulocytes and monocytes from the discovery cohort near the Guanylate Binding Protein (*GBP*) genes, whose expression has previously been associated with TB susceptibility and host response^10,18^. Almost all of these peaks were also detected as DA in the validation cohort (Fig. 2a). Genome-wide, ATB vs. HC peak height fold-changes were highly correlated between the granulocyte discovery and validation cohorts (*R*=0.93; Fig. 2b). Moreover, fold-change direction was also highly concordant (*P*-value<1e-300, Fisher’s exact test). Peak height fold-changes were less correlated in monocytes (*R*=0.70; Fig. 2b), most likely due to the smaller number of monocyte ATB samples in the validation cohort (Fig. 1b). Nevertheless, the concordance of fold-change direction was still highly significant (*P*-value=5.2e-61, Fisher’s exact test), thus confirming the robustness of DA signals detected in the discovery cohort.

**Fig. 2.**
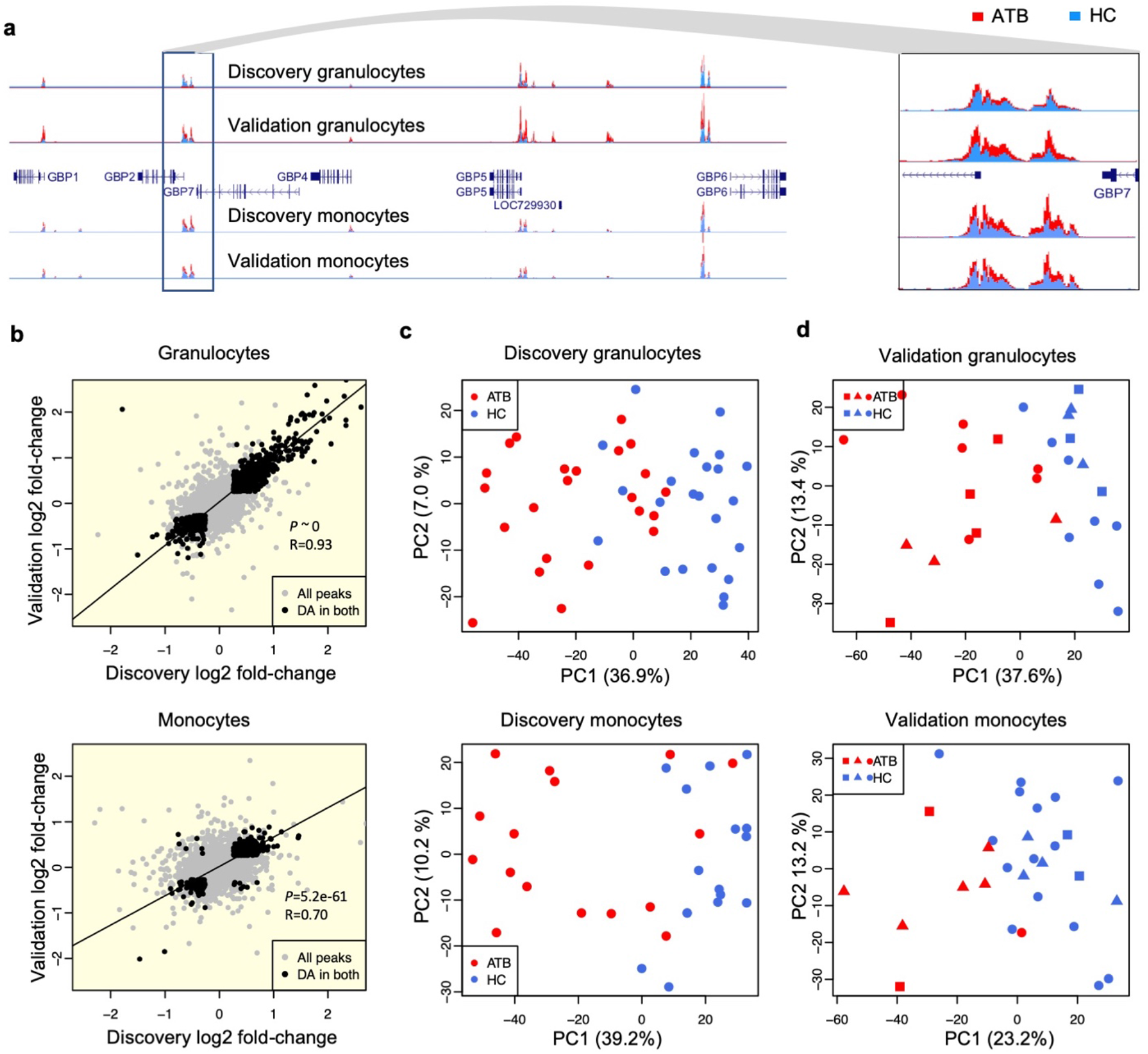
Differentially acetylated (DA) peaks between ATB and HC. (**a**) Histone acetylation profiles showing consistent differences between ATB (red) and HC (blue) in the *GBP* locus. (**b**) Scatterplot of ATB-vs-HC fold-change in discovery and validation cohorts. R indicates Pearson correlation coefficient for peaks that are DA in both discovery and validation. *P*-value of concordance in fold-change direction for shared DA peaks was calculated using Fisher’s exact test. (**c**) Discovery cohort (exclusively Chinese): principal component analysis (PCA) of granulocyte and monocyte peak heights. (**d**) Validation cohort (multi-ethnic): PCA of granulocyte and monocyte peak heights. In both (c) and (d), PCA was based on the discovery DA peak set. Squares: Indian; circles: Chinese; triangles: Malay.

To evaluate the generality of ATB-associated chromatin changes identified in the exclusively Chinese discovery cohort, we split the validation cohort into Chinese, Malay and Indian subgroups. The smaller size of these three would tend to decrease the accuracy of ATB *vs*. HC fold-change estimates, and thus reduce the correlation between discovery and validation. Nevertheless, for granulocytes, the correlation between discovery and each of the three validation subgroups remained high (*R*=0.82-0.88; Extended Data Fig. 2c). Monocytes showed a similar trend, except for the Chinese monocyte validation dataset, which contained only one ATB individual (Extended Data Fig. 2c). Thus, the global host chromatin response detected in the Chinese discovery cohort was also shared by Chinese, Malay and Indian subjects from the validation cohort.

To evaluate overall chromatin divergence between ATB and HC, we performed principal component analysis (PCA) on discovery DA peaks (Fig. 2c). As expected, the first principal component (PC1) separated ATB and HC samples in both granulocytes (*P*-value=2.2e-9; one-sided *t*-test) and monocytes (*P*-value=6.3e-6). We then used discovery DA peaks to evaluate chromatin divergence between ATB and HC in the validation cohorts. Notably, PC1 Again separated ATB from HC (Fig. 2d; granulocytes: *P*-value=1.5e-6; monocytes: *P*-value=4.8e-4), further underscoring the validity of DA peaks detected in the discovery cohort. In summary, the above results indicate that over 1,800 histone acetylation foci are altered in circulating granulocytes and monocytes in response to *Mtb* infection.

### Enriched pathways and gene loci

We used the GREAT tool^19^ to identify pathways and gene categories significantly altered in response to *Mtb* infection, based on DA peak enrichment at the corresponding gene loci. Upregulated granulocyte DA peaks were enriched for ontologies such as defense response to virus and innate immune response (Fig. 3a and Supplementary Table 2). These categories contained several type I interferon induced genes from the *IFIT* and *IFITM* families, regulators of type I interferon response (*STAT1, STAT2*) as well as interferon response factors (*IRFs*; Supplementary Table 2). Type I interferon signaling genes were also enriched, though at a lower level, near monocyte upregulated DA peaks.

**Fig. 3.**
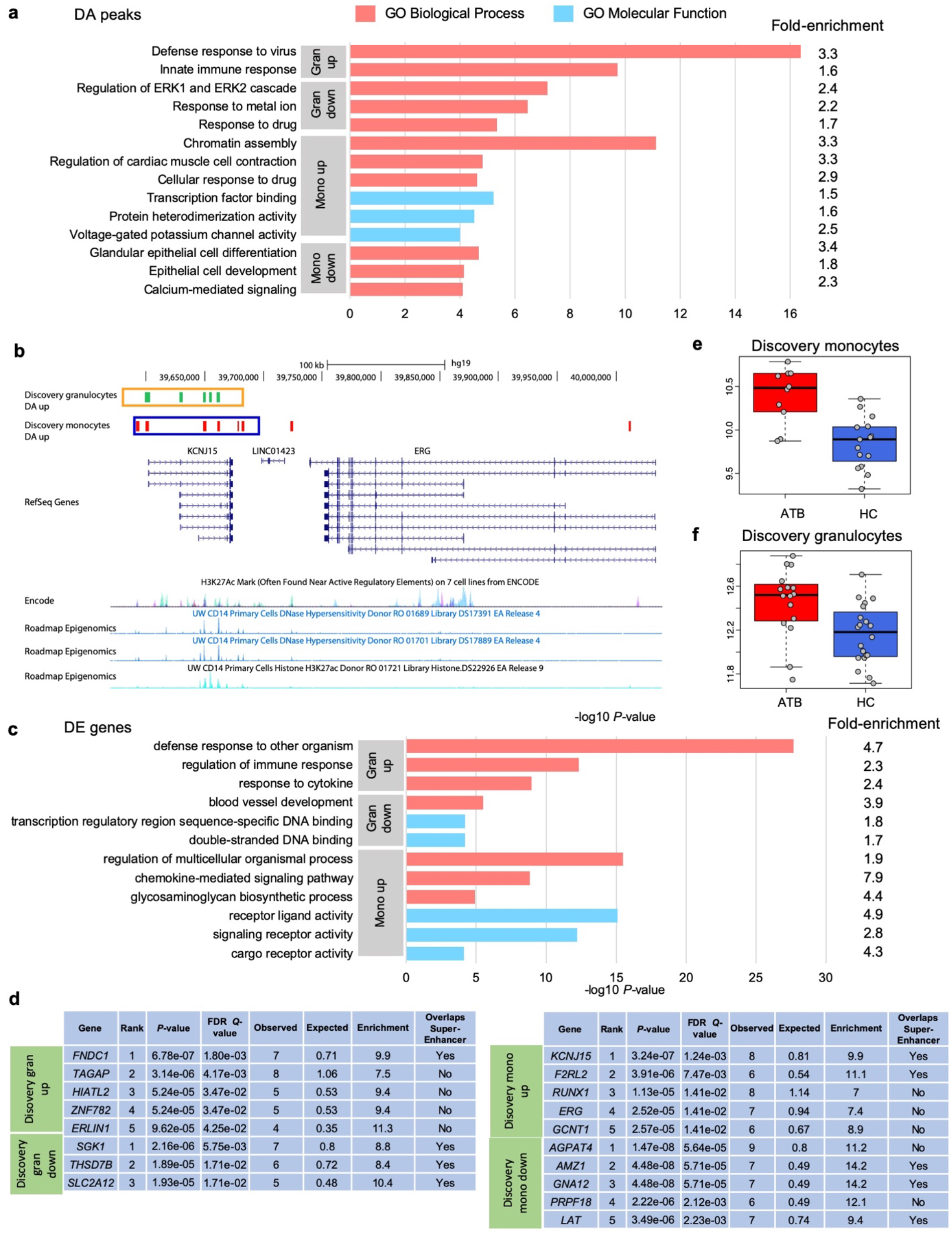
Functional enrichment analysis of DA peaks, KCNJ15 UCSC browser view and SE peak heights. (**a**) Discovery cohort: gene ontology enrichment analysis of granulocyte and monocyte DA peaks, relative to all peaks. Top three non-redundant gene ontologies (by *P*-value) are shown. (**b**) The red are monocyte DA peaks that were up regulated peaks in response to infection. The blue box indicates the monocyte super enhancer region in the *KCNJ15* locus. Similarly, green regions and the orange box indicate granulocyte DA peaks and granulocyte super enhancer region. For reference, DNaseI hypersensitivity data from CD14 primary cells (Roadmap Epigenomics), H3K27ac ChIP-seq data from CD14 primary cells (Roadmap Epigenomics) and layered H3K27ac from 7 cell types (ENCODE) are displayed. (**c**) Gene ontology enrichment analysis of DE genes, similar to (a). (**d**) Enrichment of DA peaks (relative to all peaks) near individual genes. Most significantly enriched genes are shown for each DA peak set. The last column in each table indicates whether the DA peak cluster overlaps an SE by at least one peak. (**e**) Super enhancer (SE) peak-heights of monocyte discovery samples at the *KCNJ15* locus. The peak heights were calculated from reads that mapped into the monocyte SE region displayed in (a), and covariates were regressed using the same method used for the RNA-seq data (Methods). The box plot shows that the super enhancer is activated in response to TB infection (Wilcoxson-test *P*-value=4.5e-4, fold-change=1.5; center line, median; box limits, upper and lower quartiles; whiskers, 1.5x inter quartile range; points, outliers). (**f**) Similar to (e), granulocyte samples (Wilcoxson-test *P*-value=1.7e-3, fold-change=1.3).

Upregulated monocyte DA peaks were most prominently enriched near chromatin assembly genes, indicating widespread chromatin reorganization in monocytes in response to TB (Fig. 3a and Supplementary Table 2). Notably, genes related to voltage gated potassium channel activity were also enriched (*P*-value=9.8e-5). Most prominently, the inwardly rectifying potassium channel subfamily J member 15 (*KCNJ15*) gene locus contained 8 upregulated peaks in monocytes, suggesting extensive chromatin changes upon *Mtb* infection (Fig. 3b). This locus also contained 5 upregulated peaks in granulocytes.

Next, we used GOrilla^20^ to detect functional categories enriched in DE genes (fold-change≥1.5, FDR *Q*-value≤0.05). Genes upregulated in granulocytes showed greatest enrichment for “defense response to other organism” (*P*-value=2.2e-28; Fig. 3c), including genes associated with viral defence and innate immune response (Supplementary Table 2). The latter two functional categories were also strongly enriched for DA peaks upregulated in granulocytes (Fig. 3a). These results are consistent with gene expression changes previously observed in whole blood from ATB patients, and also with the known role of type I interferon in host response to *Mtb*^4–6,10,21^. Upregulated genes in monocytes were enriched for “regulation of multicellular organismal process” (*P*-value=3.4e-16; Fig. 3c), which included genes associated with inflammatory response and regulation of response to external stimulus (Supplementary Table 2).

To identify the most extensively altered chromatin loci in TB, we tested whether the peaks near individual genes were enriched for differential acetylation (Fig. 3d). *FNDC1* and *TAGAP*, two neighboring genes involved in G-protein signalling, showed the greatest enrichment for peaks upregulated in granulocytes. Thus, this locus, which is known for association with non-infectious inflammatory diseases such as Crohn’s disease and celiac disease^22^, may also participate in the inflammatory response of granulocytes during infection. Among genes with greatest enrichment for downregulated peaks in granulocytes, *SGK1* was the most significantly altered. This gene plays a role in granulocyte apoptosis, and its downregulation leads to a prolonged state of inflammation^23^. *AGPAT4*, the gene most significantly enriched in monocyte downregulated peaks, transfers acyl chains in diacylglycerol biosynthesis, thus making them available to further use. Intriguingly, *Mtb* utilizes host acyl chains when it acquires a dormancy-like state in macrophages^24^, a condition important for *Mtb* survival. The gene most significantly associated with monocyte upregulated peaks was the above-mentioned potassium channel gene *KCNJ15.*

We then used Homer^25^ to scan for super-enhancers (SEs)^26^ in ChIP-seq profiles from the discovery cohort. SEs are known to strongly activate gene expression in a cell-type-specific manner. They are thus likely to be important for the functioning of the corresponding cell type, though it is not clear if they are functionally distinct from regular enhancers that drive strong gene expression^27^. We identified 757 SEs in granulocytes and 905 in monocytes (Supplementary Table 1). Notably, many of the DA peak clusters overlapped SEs (Fig. 3d), which further supports the significance of the corresponding DA loci in host response to ATB. In particular, the upregulated monocyte DA peak cluster near *KCNJ15* was mostly contained within an SE that also showed significantly increased acetylation in ATB (Fig. 3b,e,f).

### Role of *KCNJ15* in host response to TB

We prioritized *KCNJ15* for functional validation because: 1) this locus showed the greatest enrichment for upregulated monocyte DA peaks, 2) the DA peaks coincided with an SE, 3) its role in monocyte physiology, infection and inflammation has not been previously studied and 4) potassium channels have not been studied in the context of mycobacterial infection. *KCNJ15* encodes the poorly characterized two-transmembrane tetrameric inward-rectifier type potassium (K+) channel Kir4.2^28–30^, which potentially allows K^+^ to flow into cells when hyperpolarized^31^. Consistently, patch-clamp analysis in the human monocyte cell line THP-1 revealed an inwardly rectifying K^+^ current that was blocked by Ba^2+^ in a voltage-dependent fashion (Extended Data Fig. 3). To corroborate the increased histone acetylation at *KCNJ15* in granulocytes and monocytes (Fig. 3b,d,e,f), we analyzed *KCNJ15* mRNA expression in five different TB cohorts (Supplementary Table 3). In all five cohorts, *KCNJ15* mRNA levels in granulocytes, monocytes or whole blood were significantly higher in ATB than in HC, with monocytes showing the greatest change (Fig. 4a and Extended Data Fig. 4a,b). In the UK dataset, *KCNJ15* expression in whole blood was attenuated after two months of treatment and reached levels comparable to those of healthy individuals within 12 months (Extended Data Fig. 4b). *KCNJ15* expression was also found to be significantly upregulated in whole blood from Thai ATB patients^32^. Despite the consistent evidence from transcriptomics studies, *KCNJ15* has not previously been prioritized for mechanistic analysis of host response to TB, perhaps because other genes showed even greater differential expression. In terms of histone acetylation response, however, this locus is a clear outlier and thus a strong candidate for functional studies.

**Fig. 4.**
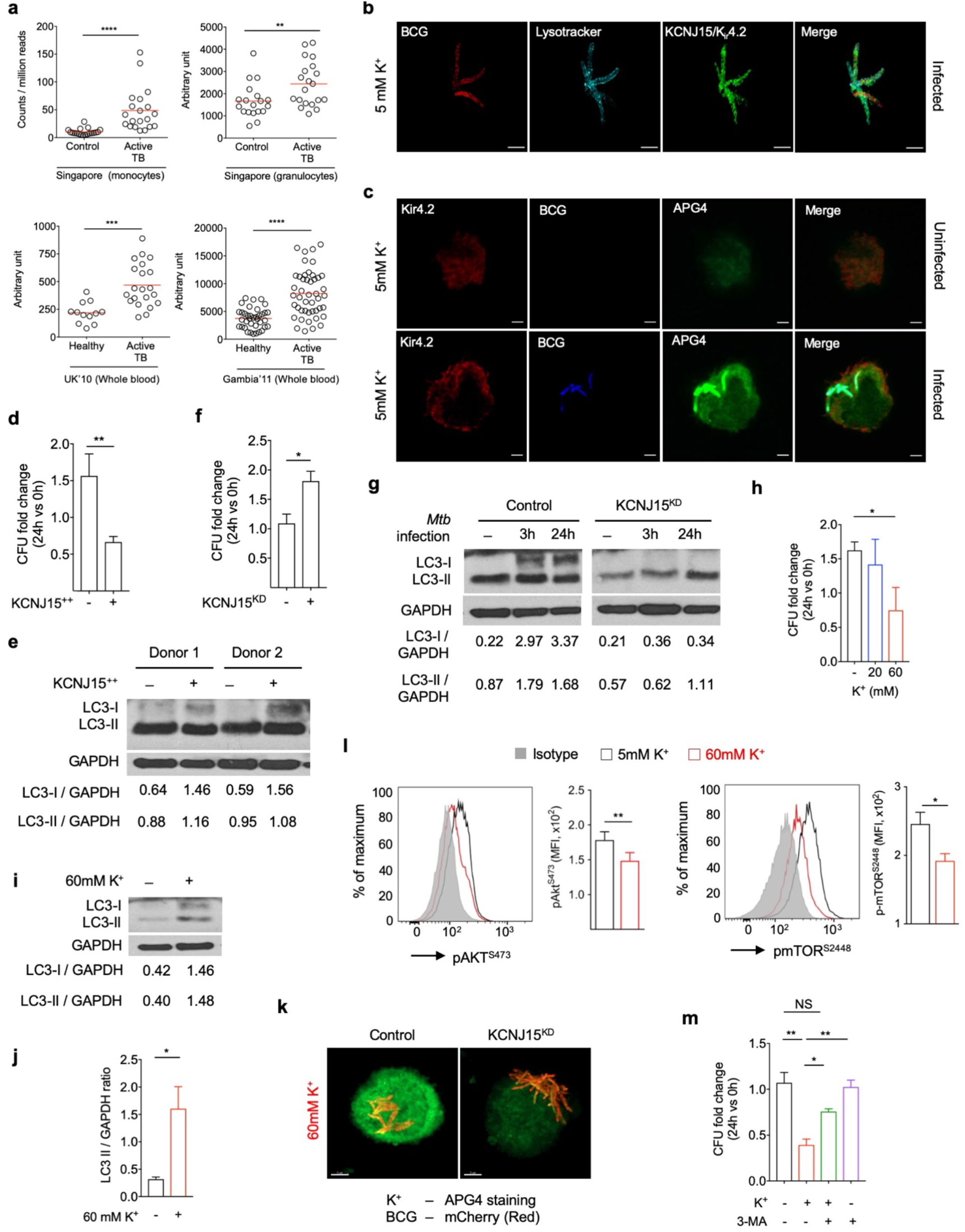
Functional characterization of *KCNJ15*/Kir4.2. (**a**) *KCNJ15* mRNA expression in ATB and HC individuals from multiple cohorts. Singapore: RNA-seq, this study; UK 2010, Gambia 2011: microarray. Red line: median. *P*-value: Mann-Whitney *U* test. (**b**) mcherry-BCG infected THP-1 monocytes stained for lysosomes and *KCNJ1*5/Kir4.2, 24hr post-infection. (**c**) Uninfected and mcherry-BCG infected THP-1 monocytes stained for intracellular K^+^ (APG-4) and Kir4.2, 24hr post-infection. (**d**) *Mtb* growth in control and KCNJ15-overexpressing (*KCNJ15*^++^) primary monocytes; n=5 donors, 24hr post-infection. CFU: colony-forming units. (**e**) Western blot, LC3 (autophagy marker) and GAPDH in *KCNJ15*^++^ and control primary monocytes, n=2 donors. Densitometric quantification of LC3/GAPDH ratio is shown below. (**f**) *Mtb* growth after 24hr in scrambled control (-) and *KCNJ15^KD^* (+) THP-1 monocytes; n=3 independent experiments, each in triplicate. (**g**) Similar to (e), *KCNJ15^KD^* and WT THP-1 cells 3hr and 24hr post-infection. (**h**) Effect of extracellular K^+^ on *Mtb* CFU in primary monocytes; n=4 donors. *P*-value: one-way ANOVA. (**i**) Similar to (e), primary monocytes incubated with 60mM K^+^ for 4hr. (**j**) Similar to (i), n=4 donors. (**k**) mcherry-BCG infected control and KCNJ15^KD^ THP-1 monocytes stained for K^+^, 24hr post-infection. (**l**) Flow cytometry: phosphorylated AKT^S473^ (pAKT^S473^) and pmTOR^S2448^ in primary monocytes stimulated as in (i); n=5 donors. (**m**) Effect of 60mM extracellular K^+^ and 3-methyl adenine (3-MA; autophagy inhibitor) in primary monocytes; n=5 donors. *P*-values: one-way ANOVA. Scale-bars: 2μm. All *P*-values were calculated using unpaired t-test, unless otherwise specified. *: *P*-value≤0.05; **: *P*-value≤0.01; ***: *P*-value≤0.001; ****: *P*-value ≤0.0001. Error bars: standard error.

To investigate the causal relationship between mycobacterial infection and *KCNJ15* expression, we infected primary monocytes and THP-1 cells with *Mtb* and BCG, respectively, and observed increased mRNA and protein expression upon infection (Extended Data Fig. 4c-e). Interestingly, in THP-1 cells Kir4.2 was localized to the cell membrane and BCG-containing lysosomes (Fig. 4b,c, and Supplementary Video 1). Lysosomal localization of Kir4.2 may be attributable to its three lysosomal-targeting dileucine motifs (^44^DGIYLL^49^, ^244^ESPFLI^249^ and ^364^ELRTLL^369^). In uninfected cells, Kir4.2 showed low expression and low APG-4 fluorescence (indicator of intracellular potassium concentration [K^+^]_i_; Fig. 4c). In contrast, BCG-infected cells showed higher Kir4.2 expression and intense and concentrated APG-4 fluorescence in phagosomes. These results suggest that K^+^ increases in phagosomes upon BCG infection, potentially due to Kir4.2 upregulation and localization to the organelle.

To assess the impact of KCNJ15/Kir4.2 on *Mtb* growth we performed gain and loss of function experiments. Lentivirally-mediated overexpression of Kir4.2 in primary monocytes (KCNJ15^++^; Extended Data Fig. 5a,b) resulted in *Mtb* growth inhibition (Fig. 4d). Notably, KCNJ15^++^ monocytes also showed increased expression of microtubule-associated protein 1A/1B-light chain 3 (LC3) (Fig. 4e). LC3 mediates, and has been used as a marker for, autophagy, an innate cellular mechanism for controlling intracellular *Mtb* growth^33^. Consistently, monocytes in which *KCNJ15* was knocked down (KCNJ15^KD^; Extended Data Fig. 5c,d) were less efficient at controlling *Mtb* growth (Fig. 4f) and impaired in LC3 expression (Fig. 4g). Furthermore, mycobacterial growth was restricted in a THP-1 cell line stably overexpressing *KCNJ15* (KCNJ15^OE^, Extended Data Fig. 5e). These results suggest a role for Kir4.2 in autophagy and control of intracellular *Mtb* growth.

To determine whether an increase in the intracellular K^+^ concentration ([K^+^]_i_) inhibited *Mtb* growth in THP-1, we increased the concentration of extracellular K^+^ from the normal level to that observed in necrotic lesions^34^, which are commonly observed in human TB (Extended Data Fig. 5f,g). [K^+^]_i_ rose from 131mM to 165mM as a result (Extended Data Fig. 5h). This increase in [K^+^]_i_ was similar to that observed in T cells treated in a similar manner^34^. Increased [K^+^]_i_ coincided with reduced *Mtb* growth in monocytes (Fig. 4h), increased levels of LC3 and occurrence of autophagic flux, indicating enhanced autophagy (Fig. 4i,j, and Extended Data Fig. S6a). K^+^-mediated mycobacterial growth restriction was also observed in PMA-differentiated THP-1 macrophages (Extended Data Fig. 5i). Importantly, BCG-infected KCNJ15^KD^ cells showed reduced APG4 staining when exposed to a high extracellular K^+^ concentration, indicating a role for Kir4.2 in potassium influx (Fig. 4k).

Autophagy is induced upon dephosphorylation of mTOR, which inhibits mTOR’s activity^35^. In our experiments, elevated extracellular K^+^ caused hypophosphorylation of Akt, mTOR and pS6 (an mTOR substrate), indicating decreased Akt-mTOR signaling (Fig. 4l and Extended Data Fig. 6b-d). Furthermore, suppression of autophagy by 3-methyladenine (3-MA) attenuated potassium-mediated inhibition of mycobacterial growth (Fig. 4m). These results indicate that increased [K^+^]_i_ restricts *Mtb* growth by inhibiting Akt-mTOR signaling, leading to increased autophagy. Similar suppression of Akt-mTOR signaling by increased [K^+^]_i_ has been described in T lymphocytes^36^. Overall, our results indicate that Kir4.2 is a conduit in the cell membrane for K^+^ to enter monocytes, and its upregulation is a protective host response to *Mtb* infection.

### Identification of haQTLs and association with eQTLs and GWAS

In addition to identifying disease-associated chromatin changes, HAWAS data can be used in conjunction with the G-SCI test^37^ to sensitively detect SNPs associated with population variation in histone acetylation, i.e. haQTLs^14^. These haQTLs could then be used to identify candidate causal variants in GWAS loci^14,17,37–41^. We identified 3,066 haQTLs in granulocytes and 3,377 in monocytes (FDR *Q*-value≤0.05; Fig. 5a,b, and Supplementary Table 4). Note that disease status was regressed out prior to QTL analysis (Methods), and thus the resulting haQTLs are not specific to TB. The haQTLs from our Chinese cohort were strongly enriched for high linkage disequilibrium (LD) with haQTLs previously detected in granulocytes and monocytes from a European cohort^17^. Nevertheless, >75% of our haQTLs were novel (Fig. 5c,d). Our haQTLs colocalized with eQTLs previously detected in granulocytes and monocytes^17,42,43^, indicating they are likely to contribute to gene expression variation (Extended Data Fig. 7 and Supplementary Table 5).

**Fig. 5.**
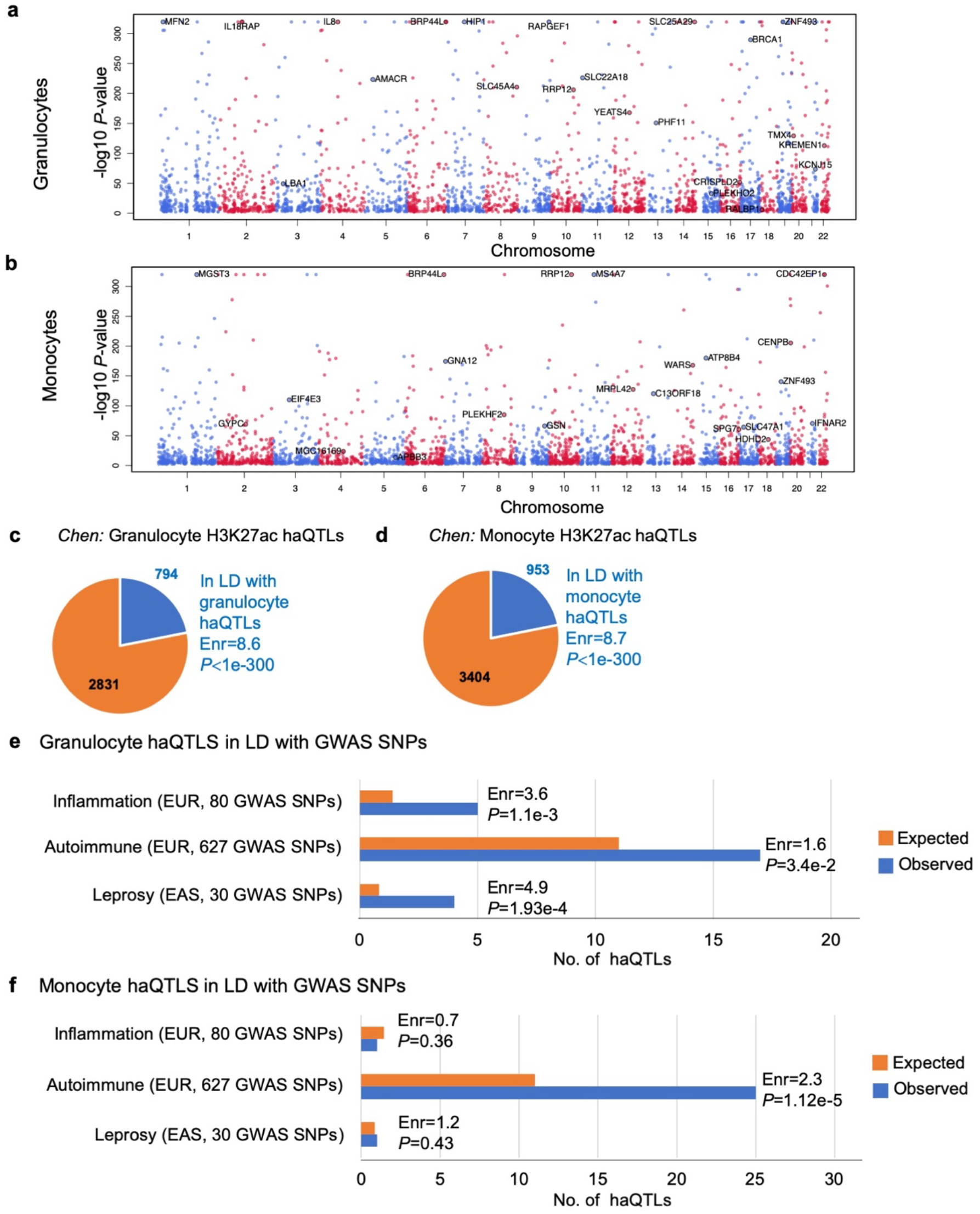
Landscape of haQTLs in granulocytes and monocytes. (**a**) and (**b**) Manhattan plots of haQTL distribution in granulocytes and monocytes. Gene names are indicated next to haQTLs that are in LD with eQTLs in the same cell type. Granulocyte eQTLs were obtained from Naranbhai et al. and monocyte eQTLs from Fairfax et al. (**c**) and (**d**) Pie charts of granulocyte and monocyte haQTLs from a previous study that are in LD with our haQTLs. (**e**)and (**f**) Number of non-redundant granulocyte and monocyte haQTLs in LD with genome-wide significant GWAS SNPs. Orange bars: expected number of haQTLs, adjusted for minor allelic frequency distribution and distance to the nearest transcription start site; blue bars: observed; Enr: fold-enrichment; *P*-value: *Z*-score test; EUR: European; EAS: East Asian.

To identify haQTLs that may contribute to human phenotypes, we tested them for LD with noncoding GWAS variants (*P*-value≤5e-8) for infectious diseases, inflammation and autoimmune disorders^44^ (Methods; Supplementary Table 6). None of the 13 TB-associated SNPs from the GWAS catalog were in LD with our haQTLs, perhaps because 10/13 were detected in non-Asian populations^45^ (Supplementary Table 6). We then examined the 30 Asian GWAS SNPs associated with leprosy, which is caused by *Mycobacterium leprae.* These 30 SNPs were enriched for LD with granulocyte haQTLs (*P*-value=1.9e-4; Fig. 5e and Supplementary Table 6), suggesting that some of the granulocyte haQTLs may contribute to mycobacterial infection by altering gene regulation in host cells^4^. We also tested the haQTL sets for association with GWAS SNPs for inflammation and autoimmune disorders (Supplementary Table 6). Most prominently, monocyte haQTLs were enriched for LD with autoimmune GWAS SNPs (*P*-value=1.1e-5; Fig. 5f). Granulocyte haQTLs showed a similar, though lower, enrichment. In addition, granulocyte haQTLs were significantly enriched in GWAS loci related to inflammation (*P*-value=1.1e-3). Thus, the haQTL sets likely contain multiple genetic variants contributing to inflammation and autoimmune disorders.

## Discussion

Our study broadens the scope of the approach by demonstrating its applicability to an immune phenotype. Since previous HAWAS analyses were based on a single cohort, the reproducibility of the detected DA peaks could not be established. Our comparison of discovery and validation cohorts demonstrates that DA peaks are indeed reproducible, despite potential confounders such as age, sex, ethnic diversity and technical variation. Importantly, this result was obtained with a relatively small cohort size, of the order of 20 disease vs. 20 control samples. The recently concluded HAWAS on Alzheimer’s disease^16^ was also based on a similarly small cohort. Thus, HAWAS is an inexpensive and robust technique, which could potentially offer mechanistic insights into diverse diseases.

In addition to identifying disease-associated chromatin traits (DA peaks), HAWAS provides a rich resource of haQTLs, some of which are likely to represent the underlying causal variants within GWAS loci. GWAS results can be population-specific, most commonly because of differences in allele frequency and linkage between population groups^46^. These two factors may also contribute to population differences in haQTLs. For example only 22% of Chinese granulocyte and monocyte haQTLs were present in the corresponding European haQTL sets^17^ (Fig. 5C). Moreover, the causal candidates inferred through LD with GWAS variants were also largely unique (Supplementary Table 6). Overall, our results suggest that haQTL analysis in novel populations could substantially expand the set of known regulatory variants and thus aid in the prioritization of causal candidates in GWAS loci.

The >2,000 differentially acetylated loci identified in this HAWAS provide a large set of candidates for locus-specific mechanistic investigations into *Mtb*–induced host response. The enrichment of DA peaks near voltage-gated potassium channels is particularly intriguing. Although this gene category has not previously been examined in the context of innate immune response to mycobacterial infection, the connection to TB is plausible because potassium channels are thought to stimulate autophagy^47,48^, a known mechanism of TB control by monocytes and macrophages^33^. Indeed, we found that upregulation of *KCNJ15*/Kir4.2 induced autophagy and restricted microbial growth in monocytes, and extracellular K^+^ supplementation had similar effects. Since potassium channels are druggable, these results open up a new avenue for research into host-directed therapies for TB^49^.

## Methods

### Human subjects

HIV-negative ATB patients (based on clinical diagnosis with mycobacterial and radiographic evidence) and HC individuals (negative for IFN-γ release assay (IGRA); QuantiFERON TB gold test) were recruited at Tan Tock Seng Hospital’s Tuberculosis Control Unit (TTSH, TBCU). Additional HC individuals were recruited locally at SIgN. Patients were sampled within 7 days of anti-TB treatment initiation and excluded if they had previously received anti-TB therapy. This study has complied with all ethical regulations and was approved by the Domain Specific Review Board (DSRB) of the National Healthcare Group (#2010/00566; for collection of blood samples from TB patients and healthy individuals), Institutional Review Board of the National University of Singapore (#09-256; for collection of blood samples from healthy individuals), and SingHealth Centralised Institutional Review Board (#2017/2806; for collection of blood samples from healthy individuals). IRB #2017/2806 is the continuation of IRB #09-256. All subjects provided consent to be included in the study.

### Reagents

The following chemicals were used: KCl (Sigma-Aldrich, #P954), Bafilomycin A1 (Sigma-Aldrich, #B1793), Phorbol Myristate Acetate (PMA) (Sigma, #P1585), 3-Methyl-Adenine (3MA) (Sigma, #M9281), Paraformaldehyde (Electron Microscopy Sciences, #15710). The following antibodies were used: anti-KCNJ15 (Sigma-Aldrich, #HPA016702), anti-LC3B (Cell Signaling Technology, #3868; Novus, #NB100-2220), anti–glyceraldehyde-3-phosphate dehydrogenase (GAPDH) (14C10) (Cell Signaling Technology, #2118), anti-rabbit immunoglobulin G (IgG) horseradish peroxidase (HRP)-linked antibody (Cell Signaling Technology, #7074), Alexa Fluor 647 mouse anti-mTOR (pS2448) (BD Biosciences, #564242), V450 mouse anti-Akt (pS473) (BD Biosciences, #560858), V450 mouse anti-S6 (pS235/pS236) (BD Biosciences, #561457), mouse IgG1 kappa isotype control eFluor 450 (eBioscience, #48-4714-82) and Live/dead fixable aqua dead cell stain kit (Thermo Fisher Scientific, #L34965). ON-TARGET plus SMARTpool Human-KCNJ15 (3772) small interfering RNAs (siRNAs) (#L006245000005) and control siRNAs (#1299001) were from Dharmacon and Integrated DNA Technologies, respectively.

### THP-1 cell line

Human monocyte THP-1 cells from ATTC were maintained in Roswell Park Memorial Institute (RPMI-1640) medium (Gibco), supplemented with 10% heat-inactivated fetal bovine serum (FBS), 1% sodium pyruvate, 1% L-glutamine, 1% non-essential amino acids (Life technologies), and 1% kanamycin (Sigma-Aldrich), at 37°C in a 5% CO_2_ humidified atmosphere. In infection experiments, no antibiotic was used. Macrophage differentiation of THP-1 cells was effected by treating with 100 ng/mL phorbol myristate acetate (PMA) for 24hr. Cells were then washed and rested for 24hr prior to infection.

### Primary Monocytes and Granulocytes

Primary cells were purified from the blood of subjects recruited in this study (see below).

### Cell separation and isolation from human peripheral blood

Peripheral blood mononuclear cells (PBMCs) and granulocytes were separated using a Ficoll gradient and RBCs were lysed. From PBMCs, monocytes were isolated using CD14^+^ immunomagnetic separation beads (MACS, Miltenyi). Isolated cells were immediately fixed with 1.6% paraformaldehyde and stored at −80°C before processing for Chip-seq. Flow cytometric analysis was performed on purified cell populations using monoclonal antibodies against CD3, CD14 and CD15. Both cell populations showed purity >95% (Extended Data Fig. 8).

### ChIP-seq of isolated granulocytes and monocytes

For each ChIP-seq experiment, approximately 10 million fixed granulocytes or 3 million monocytes were thawed on ice. Cells were lysed (10mM Tris-HCl [pH 8], 0.25% Triton X-100, 10mM EDTA, 100mM NaCl, Roche 1X Complete Protease Inhibitor) and nuclei were collected and re-suspended in 300μl SDS lysis buffer (1% SDS, 1% Triton X-100, 2mM EDTA, 50mM Hepes-KOH [pH 7.5], 0.1% sodium deoxycholate, Roche 1X Complete Protease Inhibitor). Nuclei were lysed for 15 minutes, followed by sonication to fragment chromatin to an average size of 200–500bp (Bioruptor NGS, Diagenode). Protein-DNA complexes were immuno-precipitated using 3μg of H3K27ac antibody from the same lot for all ChIP experiments (Catalogue #39133; Active Motif) coupled to 50μl Protein G Dynabeads (Thermo Fisher) overnight. Beads were washed and protein-DNA complexes were eluted using 150μl of elution buffer (1% SDS, 10mM EDTA, 50mM Tris-HCl [pH 8]), followed by protease treatment and de-crosslinking overnight at 65°C. After phenol/chloroform extraction, DNA was purified by ethanol precipitation. Library preparation was performed as in ^14^ After 15 cycles of PCR using indexing primers, libraries were size selected for 300-500bp on low-melting agarose gel and 4 libraries were pooled for sequencing at 2×100bp in each lane of an Illumina HiSeq 2000 flow cell using Version 3 reagents. In total, we generated 190 H3K27ac ChIP-seq datasets from 90 granulocyte and 100 monocyte samples.

### Read alignment, peak calling and peak height normalization

Reads were mapped to the human genome (GRCh37) using BWA-0.7.5a^50^ at default parameter settings as described previously^14,37^. Reads not annotated by BWA as properly paired were discarded, as were reads with mapping quality<10. Duplicate reads (read-pairs mapping to the same genomic location) were collapsed. For each sample, ChIP-seq peaks were detected relative to input control using DFilter^13^ at a *P*-value threshold of 1e-6. For either cell type, the initial peak set was defined as the union of peaks from the entire set of discovery and validation samples from Batches 1 and 2, which included the vast majority of samples (Supplementary Table 1). Peaks wider than 8Kb were then discarded. We performed multi-sample correction for GC bias^13,37^ separately on each processing batch. We then discarded samples with low complexity as measured by the number of unique read-pairs (<15 million for Batches 1 and 2; <30 million for Batch 3). We also discarded samples with high GC bias (>4,500 peaks showing greater than two-fold GC bias in either direction).

Of the 135 retained high-quality samples (74 granulocyte, 61 monocyte), we designated those from age-matched Chinese donors in Batches 1 and 2 as belonging to the discovery cohort (Fig. 1b). The remaining samples, which were multi-ethnic (Chinese, Malay, Indian) and included a third processing batch, constituted the validation cohort. For each combination of cell type and cohort, the final consensus peak set was then defined as the set of peaks detected in at least 5 of the retained high-quality samples (Supplementary Table 2). These consensus peaks presumably indicate the locations of active enhancers and promoters in the corresponding samples. Note that only autosomal chromosomes were considered in our analyses. Peak heights within each peak set were quantile normalized to match the distribution of mean peak height.

### Differential Peak Calling

The following 7 biological and technical confounders were considered in our analysis of peak height variation across samples in the discovery cohort: age, sex, number of reads, median insert size, number of peaks, percent unique reads, and sequencing batch. We used an iterative strategy where the confounder that explained the greatest amount of variance was regressed out at each iteration (Extended Data Fig. 1b). Note that confounding factors were regressed out from the logarithm of the peak height matrix, following the multiplicative-effects model used previously^14^. The procedure was terminated when the first 5 principal components of the residual peak height matrix were uncorrelated with any of the 7 potential confounders (Spearman correlation coefficient≤0.4; Extended Data Fig. 1c). This regression strategy accounts for the non-independence of confounding variables. By this procedure, we regressed out four covariates from the monocyte dataset (batch, no. of peaks, percent unique reads and sex) and four from the granulocyte dataset (batch, no. of reads, no. of peaks, and sex). Data from the multi-ethnic validation cohort were controlled for two additional covariates: age and ethnicity (Extended Data Fig. 1c). For each cohort, differentially acetylated peaks were determined using a 2-step approach^14^. First, using all individuals in the cohort, a differential acetylation *P*-value was calculated for each peak using the 2-sample one-sided Wilcoxon test (FDR *Q*-value≤0.05; fold-change≥1.2). We then followed a procedure designed to restrict the DA peak analysis to the core set of consistent samples, i.e. samples that displayed a global histone acetylation signature consistent with their categorization as ATB or HC^14^. Briefly, we defined the pairwise distance between two samples to be the Pearson correlation coefficient of their peak height vectors (DA peaks only). For each sample, we then calculated its median distance from HC samples and its median distance from ATB samples within the same cohort. If an HC sample was closer to ATB samples, it was discarded from the core set. Similarly, ATB samples that were closer to HC were also discarded from the core set. The final core set used for DA peak calling comprised 47 ATB and 71 HC samples (Fig. 1b). The final set of DA peaks was then defined based on samples from the core set (FDR *Q*-value≤0.05; fold-change≥1.2; Supplementary Table 1). Discovery-cohort DA peaks defined in this manner were independently corroborated using the validation cohort (Fig. 2b). Peak height heatmaps and MA plots for samples from the core set are shown in Extended Data Figs. 2a and 1d.

### Functional Enrichment Analysis of DA Peaks

We used the GREAT tool^19^ to separately determine the enrichment of gene categories in up- and down-regulated peaks. Genes were associated with regulatory regions using the basal+extension association rule defined by GREAT. The statistical significance of DA-peak enrichment near genes from any particular category (relative to all peaks) was calculated using Fisher’s exact test (FDR *Q*-value≤0.05; fold-change≥1.5). Enriched functional categories containing fewer than 6 genes flanking DA peaks were then discarded. Finally, for display in Fig. 3a, we discarded any enriched gene category if it was redundant with a higher-ranked (more significant) category. Redundancy between two functional categories was defined as overlap of ≥40% in the number of genes flanking DA peaks. The complete list of GREAT results, including redundant hits, is shown in Supplementary Table 2. Enrichment analysis of DA peaks near individual genes (Fig. 3d) was performed using the same statistical method, except that in this case each peak was assigned to its two closest genes.

### Comparison with IHEC data

We downloaded bigwig and peak files corresponding to 184 granulocyte and 173 monocyte H3K27ac ChIP-seq datasets from the IHEC Data Portal^51^. We obtained the intersection of IHEC peaks with peaks in our data (26,429 granulocyte and 33,720 monocyte peaks common to both) and used the bigWigAveToBed tool to calculate peak heights in the IHEC dataset ^52^. For comparison to IHEC, we converted BAM files from this study to bigwig format and then ran bigWigAveToBed on the same peak regions. For each peak region, we correlated mean peak height (across samples) in the IHEC dataset to mean peak height in our own.

### Detection of SEs

For each monocyte and granulocyte sample from the discovery cohort, SEs were ascertained using Homer^25^. SE annotations for each cell type were merged by constructing the union set of SEs from individual samples. Elements of the union set that overlapped SEs in fewer than 5 individuals were discarded, resulting in a final set of 757 consensus SEs in granulocytes and 905 in monocytes. SE peak height in each sample was defined as the number of ChIP-seq reads mapped to the corresponding genomic region. The SE coordinates and read counts for each sample as calculated by Homer are given in Supplementary Table 1.

### RNA-seq of isolated granulocytes and monocytes

RNA-seq was performed on 39 granulocyte and 39 monocyte samples from ATB and HC individuals (Fig. 1d and Supplementary Table 1). Total RNA was extracted using the Ambion *mir*Vana™ miRNA Isolation Kit (Ambion^®^ Thermo Fisher Scientific, #AM1561) according to the manufacturer’s protocol. All human RNAs were analyzed on an Agilent Bioanalyzer for quality assessment. RNA QC: RNA Integrity Number (RIN) ranged from 7.1 to 10, with a median value of 9.5. cDNA libraries were prepared using 300 ng of total RNA and 2ul of a 3:1000 dilution of ERCC RNA Spike-In Controls (Ambion^®^ Thermo Fisher Scientific, #4456739). The fragmented mRNA samples were subjected to cDNA synthesis using Illumina TruSeq^®^ RNA sample preparation kit version 2 (Illumina, #RS-122-2001 and #RS-122-2002) according to the manufacturer’s protocol, except for the following modifications: 1. use of 12 PCR cycles, 2. two additional round of Agencourt Ampure XP SPRI beads (Beckman Courter, #A63881) to remove >600bp double-stranded cDNA. The length distribution of the cDNA libraries was monitored using DNA 1000 kits on the Agilent bioanalyzer (Agilent, Santa Clara, CA, USA). All samples were subjected to an indexed PE sequencing run of 2×51 cycles on an Illumina HiSeq 2000 (Illumina, San Diego, CA, USA) (3-4 samples/lane). FASTQ files were mapped using STAR to the human genome build hg19. The mapped reads were then counted using featureCounts based on GENCODE V19 annotations to generate gene counts.

### RNA-seq data analysis

Gene-specific read counts were normalized to RPKM values and a consensus set of expressed genes was defined as those that were expressed (log2(RPKM)≥-0.5) in at least 5 individuals. This yielded 12,465 expressed genes for granulocytes and 13,309 for monocytes. One granulocyte sample was discarded due to poor library quality (percent reads mapped=45.3% and percent uniquely mapped reads=43.1%). Age, sex, ethnicity, and four technical covariates (number of reads, fraction of reads in exonic regions, percent of reads mapped to the human genome, percent uniquely mapped reads) were regressed out from the log2(RPKM) values (Extended Data Fig. 2b). Differentially expressed (DE) genes were subsequently ascertained by the same two-step procedure as was used on ChIP-seq data (Supplementary Table 1). As before, we used the two-sample one-sided Wilcoxon test to quantify statistical significance (FDR Q-value≤0.05; fold-change≥1.5). Differential expression was correlated to differential acetylation by associating DE genes to all DA peaks within 10Kb of their transcription start site (Fig. 1f). Gene ontology analysis was performed using GOrilla^20^ provided with a foreground (DE genes) and background list (all expressed genes). The statistical significance was computed using Fisher’s exact test (FDR *Q*-value≤0.05; fold-change≥1.5) and redundant terms were defined as above (overlap of ≥40% in the number of genes). The complete list of GOrilla results including redundant terms is shown in Supplementary Table 2.

### Transcriptomic data

*KCNJ15* gene expression profiles were analyzed in the 77 RNA-seq samples (Fig. 1c) and four publicly available clinical datasets of whole blood gene expression profiles from ATB patients (n=92) and healthy individuals (n=61), patients with TB undergoing therapy at various time points (n=7) (Supplementary Table 3). The following primers were used to measure *KCNJ15* expression by quantitative RT-PCR in *in vitro* experiments: F – CCCGGTGAGCCCATTTCAAATC and R- GACCAACTGAGCAACCAACAGG.

### Quantification of intracellular potassium ([K^+^]_i_) by flow cytometry

THP-1 cells were loaded with potassium sensing dye Asante Potassium Green (APG-4) (TEFLabs) using PowerLoad (Invitrogen) according to the manufacturer’s protocols. To calibrate [K^+^]_i_, APG-4 loaded cells were treated with 50 μM amphotericin-B (to increase membrane permeability to monovalent ions) in increasing doses of extracellular K^+^: 5, 40, 80, 120, 140, 160 and 180mM, to provide a known [K^+^]_i_. [K^+^]_i_ in various extracellular K^+^ conditions were then quantified based on intra-experimental calibration curve. All experiments were carried out on a BD LSRII Fortessa flow cytometer (BD Biosciences) and data was analysed with FlowJo (Three Star).

### Confocal and 3D-SIM super-resolution microscopy

Following infection with BCG, THP-1 cells were seeded onto coverslips. mcherry-BCG infected cells were loaded with 5μM APG-4 (TEFLabs) with PowerLoad (Invitrogen) while GFP-BCG infected cells were stained with LysoTracker Red DND-99 (Molecular Probes), according to manufacturers’ instructions respectively. Cells were then fixed in 4% (v/v) formaldehyde, permeabilized with 0.3% (v/v) Triton-X 100 in PBS and blocked with 5% (w/v) BSA in PBS. Cells were stained for Kir4.2 (Sigma) followed by secondary antibody conjugated with Alexa Fluor 647 (Molecular Probes) and Hoechst 33258 (Sigma). Stained cells were mounted with Vectashield H-1000 (Vector Laboratories). Confocal imaging was performed using a laser scanning microscope LSM 800 with Airyscan (Carl Zeiss). A Plan-Apochromat 63X/1.40 Oil DIC M27 objective lens and excitation wavelengths of 405, 488, 561 and 640 nm were used. The confocal pinhole was set to 1 Airy unit for the green channel and other channels adjusted to the same optical slice thickness. Airyscan processing was performed with ZEN software platform (Carl Zeiss). 3D-SIM images were acquired on a DeltaVision OMX v4 Blaze microscope (GE Healthcare) equipped with 405, 488, 561 and 647 nm lasers and a solid-state illuminator (WF) for excitation, and a BGR filter drawer (emission wavelengths 436/31 for DAPI, 528/48 for Alexa Fluor 488, 609/37 for Alexa Fluor 568, and 683/40 for Alexa Fluor 642). An Olympus Plan Apochromat × 100/1.4 Point Spread Function (PSF) oil immersion objective lens was used with liquid-cooled Photometrics Evolve EM-CCD cameras (Photometrics, Tucson, AZ) for each channel. Fifteen images per section per channel were acquired (made up of three rotations and five phase movements of the diffraction grating) at a z-spacing of 0.125μm. Structured illumination deconvolution followed by alignment was carried out with SoftWorX (Applied Precision). Three-dimensional image reconstruction, figure and movie preparations were done with Imaris software (Andor-Bitplane, Zurich).

### Patch-clamp studies

THP-1 cells were centrifuged at 100 g for 3 min at room temperature. The supernatant was removed and cells were suspended in 1 mL of CHO serum-free medium and transferred into Qfuge tubes. Cells were washed two additional times with extracellular solution used for QPatch experiments, and then suspended in 150-350μL extracellular solution to keep cell density at 2-5×10^6^/ml. Whole-cell patch-clamp experiments were carried out on a QPatch-48 automated electrophysiology platform (Sophion Biosciences) using disposable 48-channel planar patch chip plates. Cell positioning and sealing parameters were set as follows: positioning pressure −70 mbar, minimum seal resistance 0.1 GΩ, holding potential −80 mV, holding pressure −20 mbar, the minimum seal resistance for whole-cell requirement 0.001 GΩ. Access was obtained with the following sequence: (1) suction pulses in 29 mbar increments from −250 mbar to −453 mbar; (2) a suction ramp of amplitude of −450 mbar; (3) −400 mV voltage zaps of 1 ms duration (10×). Following establishment of the whole-cell configuration, cells were held at −20 mV for 20 ms, a voltage step protocol (−150 mV ~ +90 mV) with 10 mV increment (I-V study) or a ramp protocol (ramped from −110 to +40 mV in 200 ms). The first trace was recorded 3 min later after changing the external solution, and 10 set of traces (I-V protocol) or 10 traces (ramp protocol) were recorded. For high extracellular K^+^ or high extracellular Ba^2+^ study, the concentration was changed from low to high without washing in between. The external solution was Na^+^-Ringer’s and contained (inmM): 160 NaCl, 10 HEPES, 4.5 KCl, 1 MgCl_2_, 2 CaCl_2_ (pH 7.2, 310 mOsm). The internal solution contained (in mM): 120 KCl, 10 HEPES, 1.75 MgCl_2_, 1 Na_2_ATP, 10 EGTA and 8.6 CaCl_2_.

### Generation of VSV.G pseudotyped lentiviral and transduction of human primary monocytes

HIV-1 based lentiviral vector encoding the enhanced green fluorescent protein (EGFP) driven by CMV or the constitutive EF1a promoter for the expression of gene of interest in a SIN configuration was used. Custom synthesized (IDT biotech) human *KCNJ15* cDNA was sub-cloned into the PE1A plasmid (Invitrogen, #A10462), followed by gateway transfer of *KCNJ15* into the lentiviral vector. In brief, HEK293 cells were transfected using Xfect (TakaRa, #631318) with the viral packaging construct [pMDLg/pRRE (Addgene, #12251), pRSV/REV (Addgene, #12253), pMD2.G/V-SVG (Addgene, #8454)]. Lentiviral particles were collected from the supernatant 48 to 72hr after transfection, concentrated by LentiX concentrator (Clontech, #631232), and titrated by qPCR (determination of number of transducing or infectious units per ml) on HeLa cells. Titres for control and *KCNJ15* lentivirus were 6×10^7^ TU/ml. Blood-derived CD14+ monocytes were adjusted to a concentration of 1×10^6^ cells/ml, for gene transduction, duplicate wells of a 12-well plate were seeded with 1 ml/well of the cell suspension (1 × 10^6^ cells/ml) and virus was added in the presence of 6 μg/ml polybrene (Sigma-Aldrich, #107689) (multiplicity of infection of 25). Cells were then incubated overnight with the virus and residual vector virus and non-adherent cells were removed by washing the wells with growth medium and the plate was further incubated at 37°C. Two days after the initial addition of vector virus, transduction efficiency was determined by flow cytometry.

### Transfection of siRNAs

THP-1 cells were transfected with siRNAs using the Lipofectamine2000 Kit (Invitrogen, #11668-019) according to the manufacturer’s protocol. Knockdown was confirmed by qRT-PCR and Western blotting.

### Western blot

Protein lysates from cells were obtained by lysis in radioimmunoprecipitation assay buffer (Merck Millipore, #R0278) with protease and phosphatase inhibitors (Roche, #04906837001). A Micro BCA Protein kit (Thermo Scientific, #23235) was used to measure protein levels, and equal amounts of proteins were resolved by electrophoresis on 12% Tris-HCl gels (Mini-PROTEAN TGX gels; Bio-Rad, #4561043) and transferred onto polyvinylidene difluoride membranes (Trans-Blot Turbo Transfer Pack; Bio-Rad, #1704156). Membranes were developed using the indicated primary antibody at a 1:1000 dilution and secondary antibodies at a 1:3000 dilution in blocking solution. This was followed by incubation with Chemiluminescent HRP detection reagent (Merck Millipore, #WBKLS0500) for 1 min before image acquisition.

### Analysis of phosphorylated proteins using flow cytometry

Single-cell suspensions were resuspended in PBS and stained for 30min with live/dead fixable aqua dead cell stain kit (Thermo Fisher Scientific, #L34965). Cells were washed with FACS buffer containing PBS with 3% FBS, 1mM EDTA and 0.1% sodium azide) and incubated with Human TruStain FcX™ (BioLegend, #422302) for 20min. Following this, cells were washed with FACS buffer, fixed and permeabilized using the PerFix EXPOSE kit (Beckman Coulter, #B26976) according to manufacturer’s instructions. Permeabilized cells were incubated for 30 min in the dark with fluorophore-conjugated antibodies specific for phosphorylated marker or isotype control antibodies. Cells were analysed using the LSRII flow cytometer (BD) and data was analysed with FlowJo software.

### SNP Calling

All discovery samples indicated in Fig. 1b were used for SNP calling as described previously^14,37^. Following GATK best practices, the BAM file for each ChIP-seq sample was realigned at given indel sites and base qualities were then recalibrated^53^. For each sample, GATK’s Haplotype Caller was ran in GVCF mode, and subsequently multi-sample SNP calling was performed on all samples (granulocytes plus monocytes) using GATK’s GenotypeGVCFs tool. The raw SNP call set was filtered using our custom pipeline for SNP calling from ChIP-seq data^37^. Based on criteria defined in^37^, we filtered out SNPs that matched any of the following: QD<4.185, Inbreeding Coefficient<-0.398, within a SNP cluster of more than 19 SNPs in a 100bp window, Homopolymer run≥18, and Hardy-Weinberg equilibrium binomial test *P*-value≤1e-3. We additionally imposed the criterion that a SNP should be supported by at least 5 non-reference reads across all datasets, and at least one of the datasets should have 3 or more non-reference reads. We also discarded SNPs contained within UCSC Genome Browser self-chains blocks with a normalized score>90. Based on this pipeline, we discovered 362,021 SNPs in ChIP-seq peaks, of which 247,411 were polymorphic in the granulocyte dataset and 273,753 in the monocyte dataset.

### Genotyping

All samples were genotyped using the Illumina OmniExpress v1.1 Beadarray, which assesses >700,000 representative genetic markers in a genome-wide manner. A subset of SNPs were further validated using the Sequenom MassArray primer extension platform. Both methods have been used extensively by others and us and have been found to result in >99.99% concordance^54^. Genotypes inferred were compared against genotype calls from ChIP-seq reads to identify and correct sample swaps in the ChIP-seq pipeline.

### Histone Acetylation QTLs

We used the G-SCI test to call haQTLs in granulocytes and monocytes^37^. Only high-quality samples from the ethnically homogeneous discovery cohort were used in this analysis (46 granulocyte samples, 32 monocyte samples). The G-SCI test does not require prior knowledge of genotypes. Rather, it infers genotype likelihoods from ChIP-seq read counts and then integrates over all possible genotypes to calculate the statistical significance of an haQTL. This test combines information from peak height vs. genotype correlation across samples and allelic imbalance in read count within heterozygous individuals. These two signatures of an haQTL are combined into a single *P*-value. To infer haQTLs independently of disease state, disease status was regressed out from the peak-height matrix used for DA peak calling, and the residual was used for G-SCI analysis^14^. In this study, the G-SCI test procedure was refined in two ways. Firstly, for each SNP *k*, we replaced the uniform allele frequency prior used in the original study with a SNP-specific allele frequency *f_k_*. Thus, for each SNP, we estimated four parameters by maximum-likelihood (*f_k_*, *α*, *β* and *σ*) instead of three. Secondly, we modified the method for permutation testing to convert raw *P*-values to final *P*-values. The original permutation approach^37^ randomized the data by permuting peak heights across samples for the same SNP. In addition, reference and non-reference base calls were randomly flipped with probability 0.5 at heterozygous sites. In the current study, this procedure was generalized and accelerated by randomly swapping data between ChIP-seq units. A unit was defined as the peak height and allelic counts for a SNP in a single individual. Thus the total number of units would equal the number of SNPs times the number of individuals. Units were then binned in three dimensions according to their peak height, number of reads covering the SNP and fraction of non-reference reads. Units within the same three-dimensional bin were randomly permuted, and the haQTL *P*-value was then calculated for each SNP. As before, the permutation-test *P*-value was then calculated based on the rank of the raw *P*-value relative to the *P*-values of permuted data. Again, as before, SNPs with an adjusted *P*-value of 0 after 1 million permutations were assigned an adjusted *P*-value of 5e-7. FDR correction and filtering for QTL effect size was performed as before^37^.

### Overlap between haQTLs and eQTLs

We downloaded the granulocyte eQTLs^43^ and obtained the subset that was in tight linkage with SNPs we called within granulocyte ChIP-seq peaks. The genotypes from the European population of 1000 Genomes were used in the LD calculation, with a threshold of *R*^2^≥0.8 and maximum distance of 500 Kb. As before, we corrected for LD within the eQTL set by constructing the LD network of the eQTL set and replaced each connected component in the network by the SNP with the best eQTL *P*-value^37^. This resulted in 1,257 Naranbhai eQTLs used for analysis. We downloaded the naïve monocyte eQTLs^42^ and used the same procedure to obtain a final set of 2,216 eQTLs for further analysis. We also downloaded neutrophil and monocyte eQTLs^17^ and retained those with an FDR *Q*-value≤0.05. If a gene had more than one eQTL, the one with the best *P*-value was kept and the others were discarded. Chen et al. eQTLs were further filtered based on LD as described above, resulting in 3,069 neutrophil and 3,460 monocyte eQTLs for further analysis. Note that the vast majority of granulocytes in peripheral blood are neutrophils. For each SNP from the four eQTL sets, the LD with haQTLs from the corresponding cell type was calculated using the same 1000 Genome population and LD thresholds as above. To calculate the statistical significance of the LD, we randomly chose 1.5 million SNPs as a control set, and used a method as before wherein the likelihood of linkage of the control set and eQTL set with haQTLs was calculated while adjusting for allele frequencies, genomic location biases (distance from nearest TSS of a gene), biases in LD block sizes between the eQTL SNP and control SNP, and correcting for LD within each of the four eQTL sets^37^.

### Overlap with published haQTL datasets

We downloaded neutrophil and monocyte H3K27 haQTLs^17^ and processed them in a similar manner to eQTLs. First, we filtered for FDR *Q*-value≤0.05 and retained only the most significant haQTL for each ChIP-seq peak. We then discarded haQTLs that were not in LD with any SNP within our ChIP-seq peaks (LD was defined as *R*^2^≥0.8, distance≤500kb). As above, these were further filtered to retain only one haQTL per LD cluster, resulting in 3,625 neutrophil H3K27ac QTLs and 4,357 monocyte H3K27ac QTLs. We then calculated the statistical significance of the linkage of these haQTLs with our own haQTLs using the method described above for eQTLs.

### Statistical significance of LD between haQTLs and GWAS SNPs

We downloaded phenotype-associated SNPs from the NHGRI-EBI GWAS catalog^44^ (downloaded May 2018; Supplementary Table 6). Only SNPs with genome-wide significance (*P*-value<5e-8) were retained and duplicates were removed. GWAS SNPs annotated with GRCh38 coordinates were lifted over to GRCh37. To create the set of TB-associated SNPs, GWAS results from two TB association studies not represented in the catalog were added to the list^55,56^. Since LD with haQTLs was calculated using genotypes from 1000 Genomes data, we obtained the subset of SNPs genotyped in 1000 Genomes, resulting in 3 TB-associated SNPs in the Asian population (out of 3), 10 in the European population (out of 17) and 0 in the African population (out of 5). LD between these 13 GWAS SNPs and haQTLs was calculated using 1,000 Genomes data from the population in which the GWAS study was performed (*R*^2^≥0.8, distance≤500kb). We next downloaded 30 SNPs associated with leprosy from NHGRI-EBI GWAS catalog^44^ all of which were in East Asian populations and also genotyped in 1000 Genomes. We obtained inflammation-associated SNPs from the GWAS catalogue^44^, and found that 80 of the 81 EUR SNPs were genotyped in 1000 Genomes. We also used a set of 627 EUR autoimmune GWAS SNPs^37^ with *P*-value≤5e-8 (Supplementary Table 6). To calculate the statistical significance of linkage between GWAS SNP sets and haQTLs (leprosy, autoimmune and inflammation; Fig. 5e,f), we used a method that corrects for allele frequency, distance to the nearest transcription start site and biases in LD block size between haQTL and control SNPs^37^.

## Acknowledgements

We thank the personnel of the Biological Resource Centre’s BSL3 laboratory and the Defence Science Organization’s BSL3 laboratory for facilitating this study. This project was supported by core funds from Singapore’s Agency for Science, Technology and Research (A*STAR); A*STAR translational program in infectious disease #IAF11003; A*STAR Joint Council Office Grant #JCO-CDA15302FG151; BMRC-SERC grant #1121480006; SIgN Immunomonitoring platform grant #IAF311006; BMRC transition fund grant #H16/99/b0/011; Swiss National Foundation grant #310030-173240, the European Union’s Research and Innovation Program grant #TBVAC2020 643381, and Nanyang Technological University Singapore’s Lee Kong Chian School of Medicine Start Up Grant.

## Author contributions

S.P. and A.S. conceived the idea. R.D.R., J.P., G.D.L., A.S and S.P. designed the study and analyzed the data. R.D.R., J.P., P.K., C.Y.C., C.L., J.L., B.L., F.Z., M.P., S.T.O., H.S.H., M.M., X.L, A.L., Y.T. performed the experiments. K.G.C. designed and analyzed the electrophysiological and APG-4 data. D.P. and O.R. contributed to overexpression experiments. C.B.E.C and Y.T.W provided clinical samples and interpreted results. C.C.K. designed and performed array genotyping. J.P, R.D.R., G.D.L, A.S and S.P wrote the manuscript. All authors discussed results and contributed to the manuscript.

## Data and materials availability

ChIP-seq data have been deposited at the European Genome-phenome Archive EGA, http://www.ebi.ac.uk/ega/), which is hosted by the EBI, under accession number EGAS00001002997. RNA-seq data have been deposited at NCBI’s Gene Expression Omnibus through GEO Series accession number GSE126614.

