## Supplementary Material for "Histone acetylome-wide association study of tuberculosis"

### Extended Data Figures

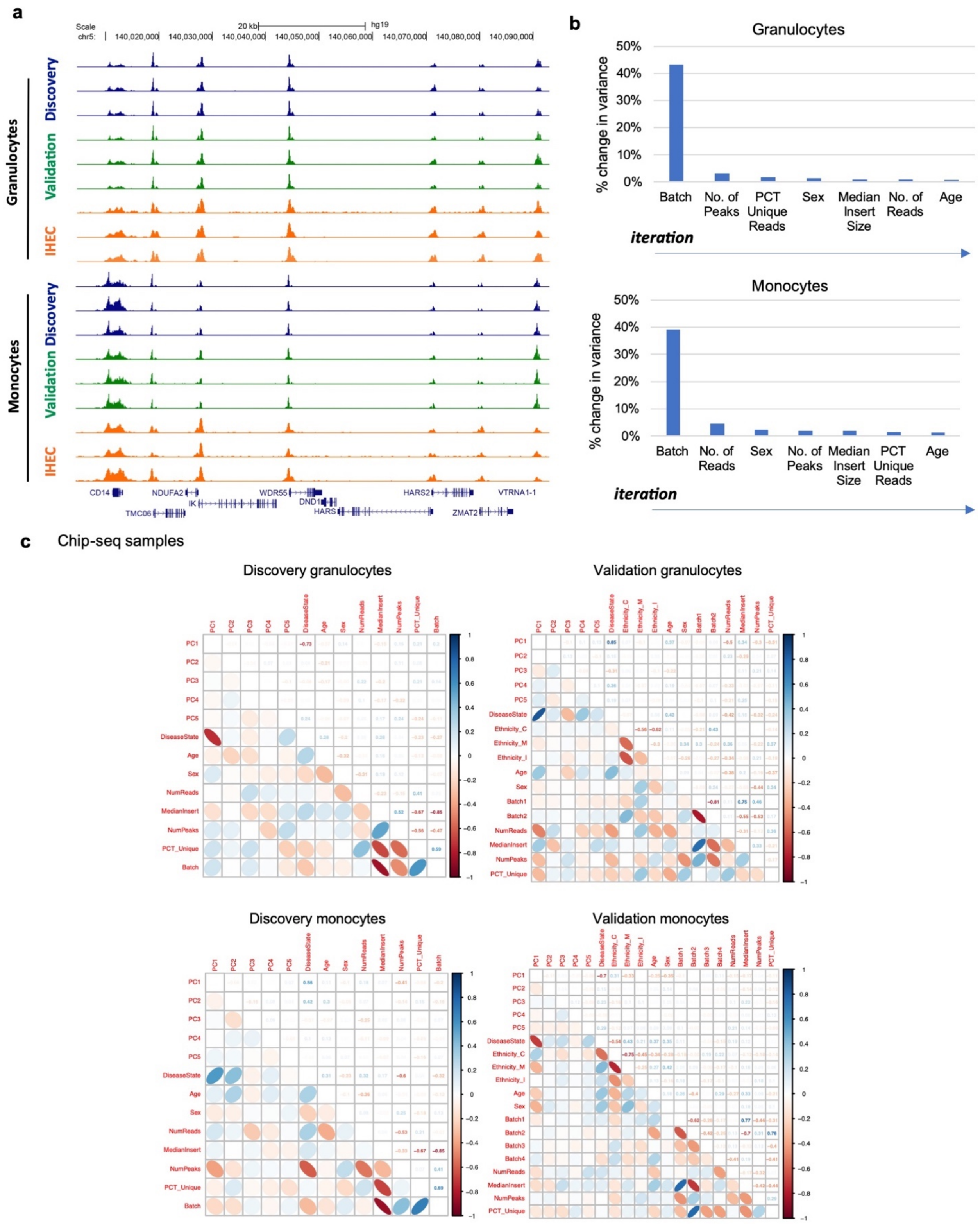

**d** Chip-seq samples

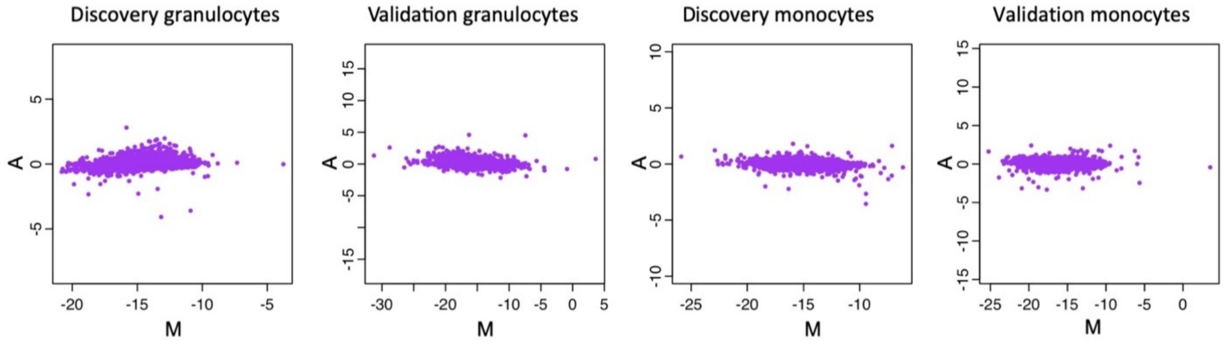


**Extended Data Fig. 1. Comparison of a locus with IHEC H3K27ac ChIP-seq and regression of confounding variables.**

(**a**) UCSC Genome Browser view of a locus: comparison of representative granulocyte and monocyte H3K27ac ChIP-seq profiles from discovery and validation cohorts with corresponding data from IHEC. Genome-wide correlation of average peak height between discovery cohort and IHEC: $R$=0.90 (granulocytes); $R$=0.87 (monocytes). (**b**) Method for selecting covariates to be regressed in discovery samples. The covariate that explained the most variance at each iteration of the method was plotted. (**c**) Correlation among the first 5 PCs of the residual peak height matrix and all biological and technical covariates. (**d**) MA plot of peak heights for the four cohorts. Only consistent samples were used.

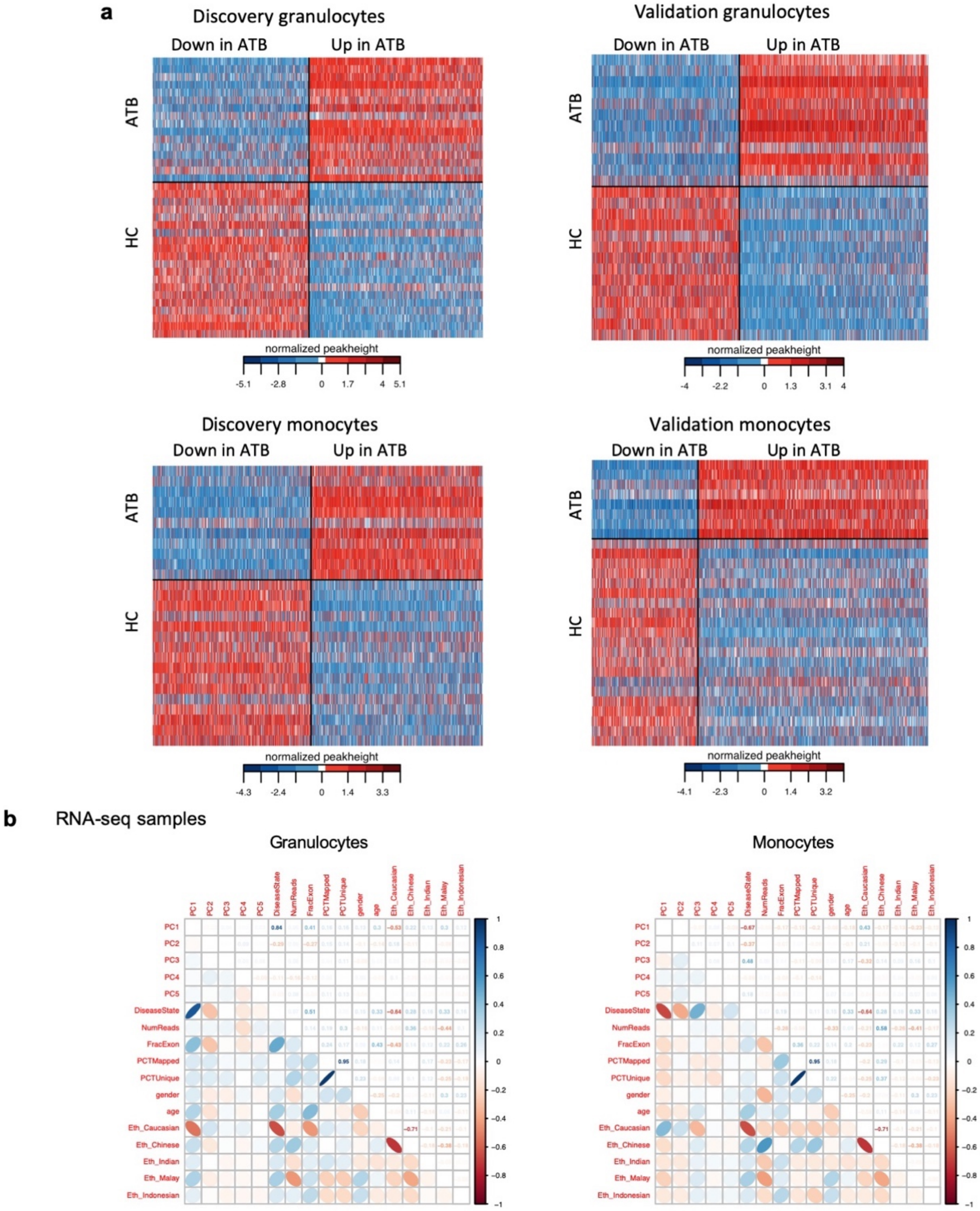
a
Discovery granulocytes
Down in ATB
Up in ATB
ATB
HC
normalized peakheight
Validation granulocytes
Discovery monocytes
Validation monocytes
b
RNA-seq samples
Granulocytes
Monocytes
PC1
PC2
PC3
PC4
PC5
DiseaseState
NumReads
FracExon
PCTMapped
PCTUnique
gender
age
Eth_Caucasian
Eth_Chinese
Eth_Indian
Eth_Malay
Eth_Indonesian

c

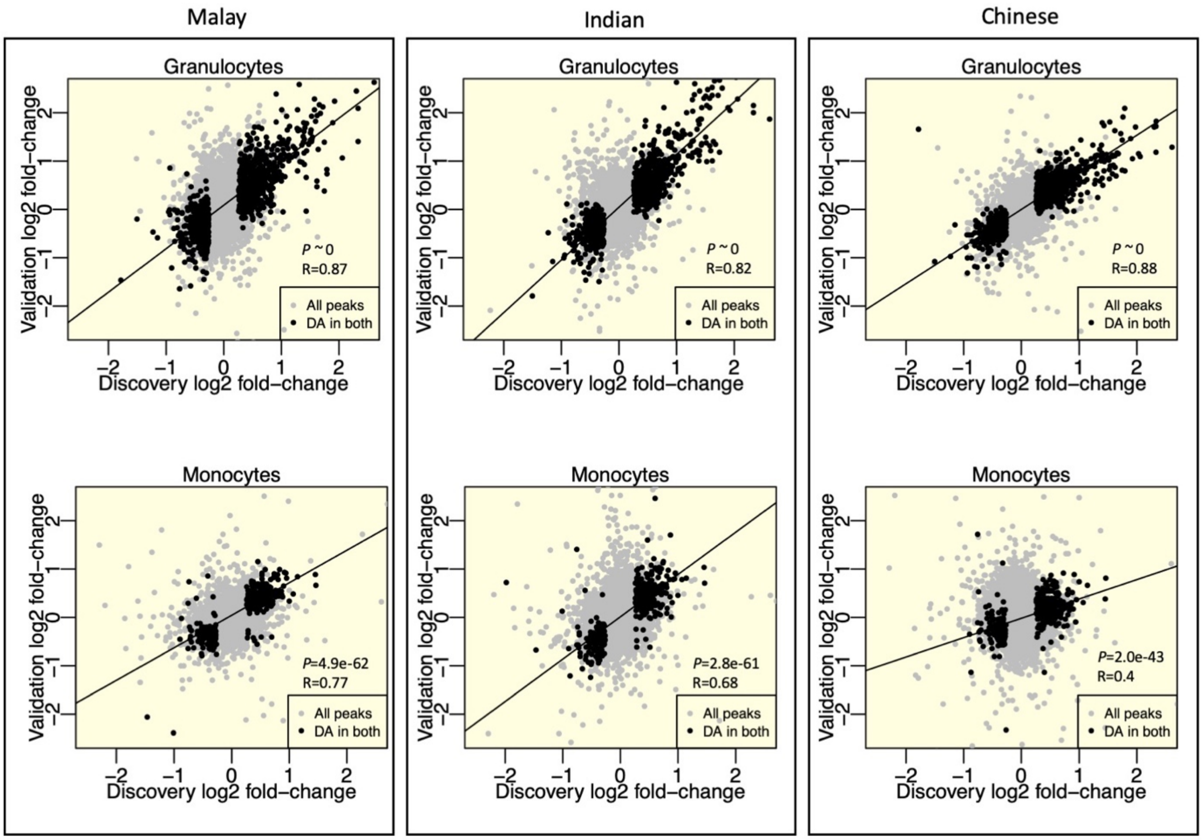


**Extended Data Fig. 2. ChIP-seq peak height z-scores, ethnicity-specific fold-change correlation and regression of confounders for RNA-seq data.**

(**a**) DA peak height z-scores for the four ATB vs. HC datasets from samples from the core set. Each row represents a single individual, and each column a DA peak. DA peaks are grouped into "down in ATB" and "up in ATB", and sorted by decreasing fold-change in each group. (**b**) Correlation among the first 5 PCs of the residual RNA-seq gene expression matrix and all biological and technical covariates. (**c**) Ethnicity-specific fold-change correlation between discovery and validation cohorts.

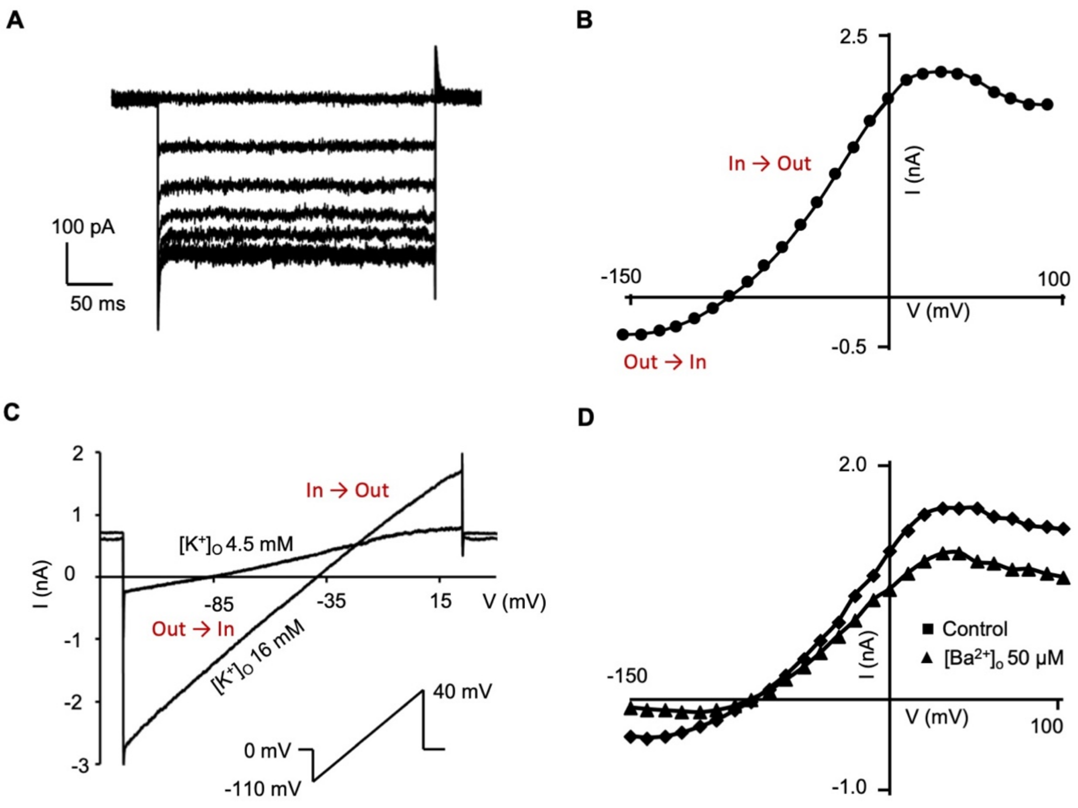


**Extended Data Fig. 3. Inwardly rectifying $K^+$ currents detected in THP-1 cells.**

**(a)** Currents elicited by stepping from a holding potential of -20 mV to -150 mV to 0 mV in 10 mV increments. The amplitude of the inward current increased with hyperpolarization. **(b)** Current - voltage (I-V) relation in a THP-1 cell. **(c)** Ramp from -110 mV to 4 mV showing the shift in reversal potential with the change of $[K^+]_o$ from 4.5mM to 16mM. **(d)** Whole-cell current - voltage (I-V) curve showing amplitudes of inwardly rectifying current, inhibited by 0.05mM external $Ba^{2+}$.

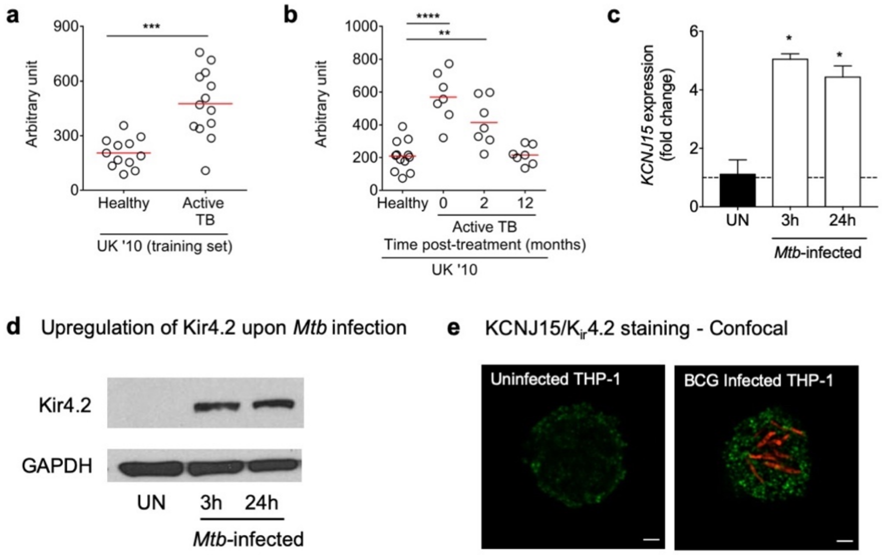


**f** The exact *P*-values for panels A-C are as follows:

| Panel | Comparison details | *P*-values |
|---|---|---|
| A | UK'10, Healthy (n=12) vs Active TB (n=13) | 0.0002 |
| B | UK'10, Healthy (n=12) vs 0 months treatment (n=7) | <0.0001 |
| | UK'10, Healthy (n=12) vs 2 months treatment (n=7) | 0.0018 |
| C | UN vs 3 h | 0.0173 |
| | UN vs 24 h | 0.0333 |

**Extended Data Fig. 4. Mycobacterial infection increases *KCNJ15*/Kir4.2 expression in whole blood, primary monocytes and THP-1 monocytes.**

(**a**) *KCNJ15* expression in the peripheral blood of UK 2010 cohort of healthy individuals and ATB patients. Higher expression of *KCNJ15* mRNA was observed in ATB patients in this cohort. Red line indicates median. (**b**) *KCNJ15* expression in the peripheral blood of ATB patients (UK 2010 cohort) during anti-TB therapy. Red line indicates median. (**c**) *KCNJ15* mRNA was assessed by qRT-PCR in *Mtb*-infected primary monocytes. *KCNJ15* expression normalized to *GAPDH*, relative to uninfected (UN) cells, is shown. Error bars: standard error. (**d**) Western blot analysis of KCNJ15/Kir4.2 and GAPDH in *Mtb*-infected primary monocytes as in (c). (**e**) Immunostaining (confocal microscopy) of KCNJ15/Kir4.2 in THP-1 cells, which was increased upon *M. bovis* BCG infection. Green: Kir4.2; red: mcherry- BCG. Scale-bars: 2μm. (**f**) *P*-values for panels a-c. **P*-value<0.05, ***P*-value<0.01, ****P*-value<0.001 *****P*-value<0.0001, by Mann-Whitney *U* test and paired Wilcoxon signed-rank test.

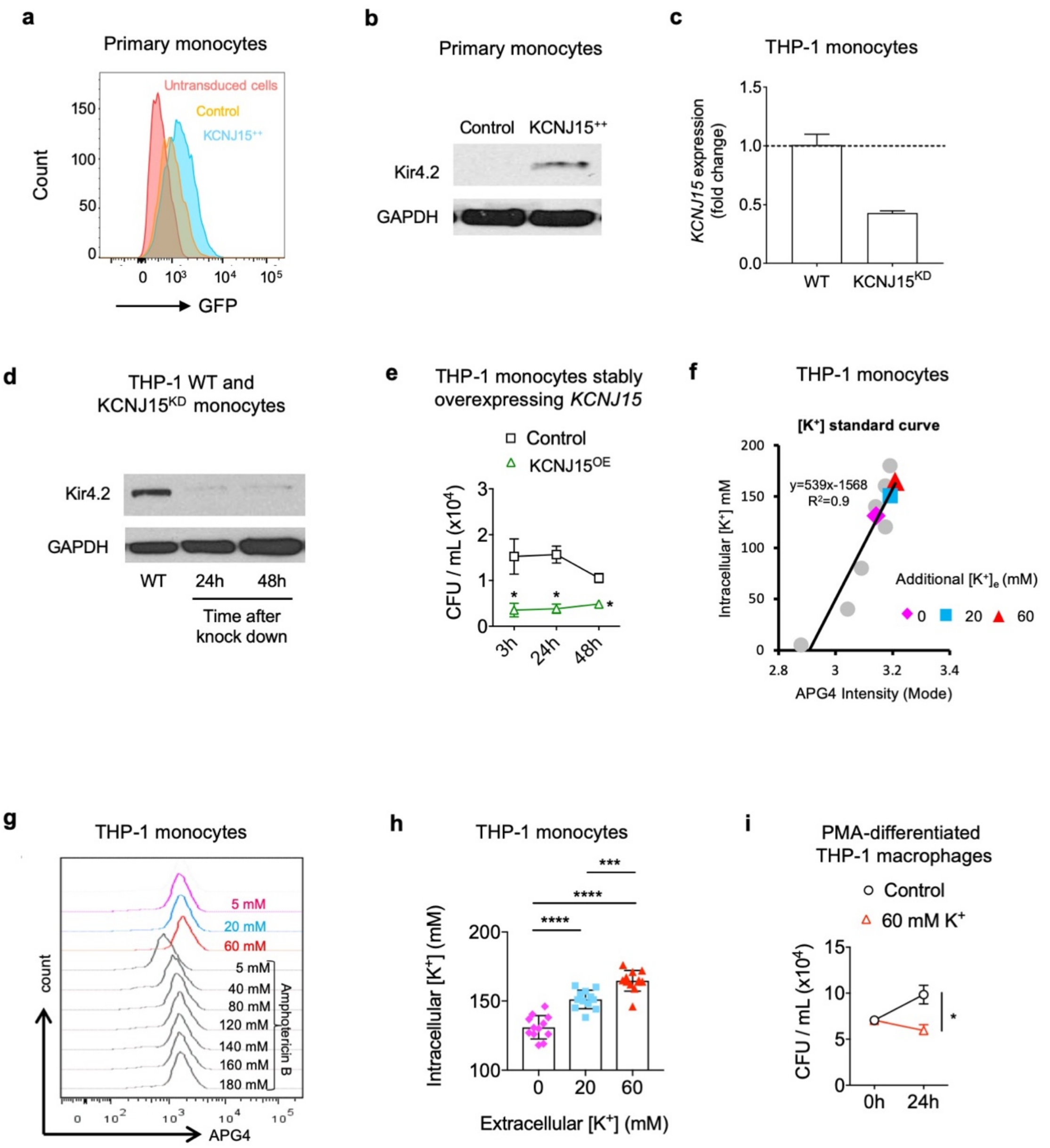


**Extended Data Fig. 5. Overexpression and Knockdown of KCNJ15/Kir4.2 in monocytes and APG-4 mediated estimation of intracellular potassium.**

(**a**) CD14$^+$ primary monocytes were transduced with either lentiviral plasmid carrying human *KCNJ15* cDNA or the control plasmid. Transduction efficiency was determined by FACS using GFP which is constitutively expressed. Pink: untransduced cells; orange: cells transduced with empty vector; blue: cells transduced with*KCNJ15* plasmid. (**b**) Transduction efficiency of primary monocytes in (a) was determined by Western blotting. (**c**) siRNA mediated abolishment of *KCNJ15* in THP-1 cells ($KCNJ15^{KD}$), RT-PCR of *KCNJ15* gene. *P*-value=0.0004, Student's *t*-test. Error bars: standard error. (**d**) Western blot for *KCNJ15*/Kir4.2 protein in knocked down ($KCNJ15^{KD}$) THP-1 cells. (**e**) Mycobacterial (BCG) growth at respective time points in control and stable THP-1 monocytes overexpressing *KCNJ15* ($KCNJ15^{OE}$) THP-1 monocytes. Two independent experiments were performed, each in triplicate. Error bars: standard error. (**f**) Calibration of intracellular $K^+$ ($[K^+]_i$) concentrations by flow cytometry in THP-1 cells. Standard curve was generated by treating APG-4 loaded cells with 50mM amphotericin-B (to increase membrane permeability to monovalent ions) in titrated doses of extracellular $K^+$ (all grey circles). Intra-experimental quantification of $[K^+]_i$ in physiological condition (5mM, magenta diamond) or high extracellular $K^+$ conditions (20 or 60mM, blue square and red triangle). The extracellular K+ concentration range was selected to match levels observed in necrotic tissue (Eil et al., 2016), since TB is characterized by necrotic TB lesions. (**g**) Histogram plots of APG-4 intensities of THP-1 cells treated with various extracellular $K^+$ conditions from (E). (**h**) $[K^+]_i$ concentrations of THP-1 cells cultured in 5mM (physiological control) and high extracellular $K^+$ medium (20mM and 60mM). N=6 (12 sets of calibrations). *****P*-value<0.0001, ****P*-value=0.0003 (1-way ANOVA, Tukey's multiple comparisons test), Baseline $[K^+]_i$ of THP-1 was quantified as 130.9±2.4mM. Extracellular $K^+$ of 20mM or 60mM increased $[K^+]_i$ to 151±1.9mM and 164.6±2.2mM respectively. Error bars: standard deviation. (**i**) Effect of 60mM K+ on mycobacterial (BCG) CFU growth after 24hr in PMA-differentiated THP-1 macrophages. Two independent experiments were performed, each in triplicate**.** Error bars: standard error.

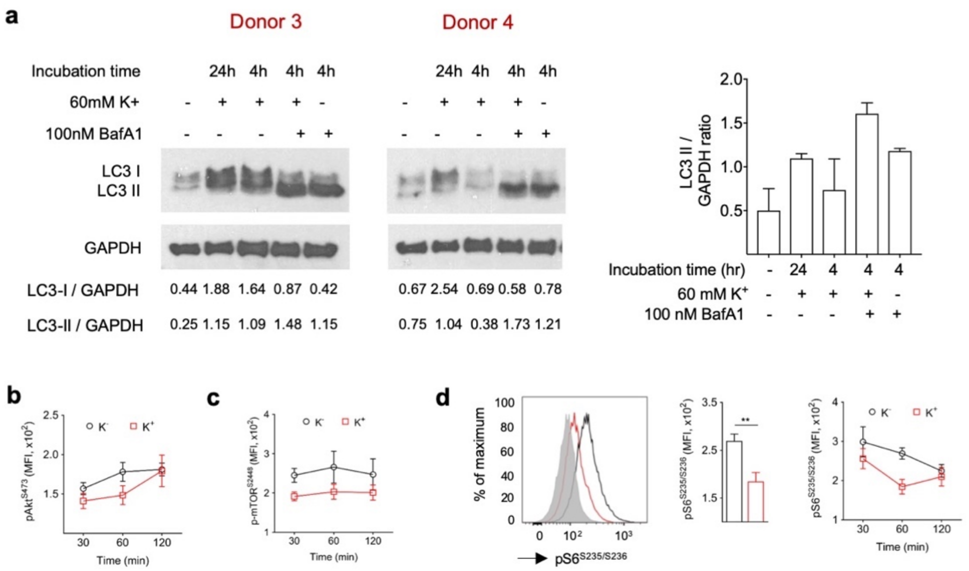


**Extended Data Fig. 6. Extracellular potassium induces autophagy and enhances AKT-mTOR signaling.**

(**a**) Western blot analysis for LC3 and GAPDH in primary CD14$^{+}$ monocytes treated with extracellular 60mM K$^{+}$ in the absence or presence of lysosomal inhibitor Bafilomycin A1 (BafA1). BafA1 increased LC3-II accumulation in monocytes treated with extracellular 60mM K$^{+}$, indicative of productive autophagosome recycling (autophagic flux). Data from two donors is shown. LC3-I/GAPDH and LC3-II/GAPDH ratios in respective conditions has been indicated. A concatenated data of LC3-II/GAPDH from different donors has been plotted (right side). (**b**) Quantitative flowcytometric analysis for phosphorylated AKT$^{S473}$ in primary monocytes treated with 60mM K$^{+.}$ Data is presented as a time course showing that K$^{+}$ suppresses Akt phosphorylation at residue S473. (**c**) and (**d**) pmTOR and pS6 analysis as in (b). Three independent experiments were performed. **, *P*-value=0.0014, Student's *t*-test. Error bars in all panels: standard error.

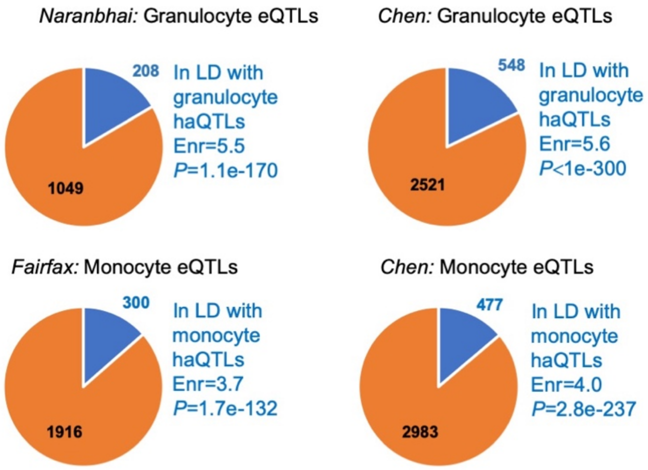


**Extended Data Fig. 7. Pie charts of granulocyte and monocyte eQTLs from previous studies that are in LD with corresponding haQTLs from this study.**

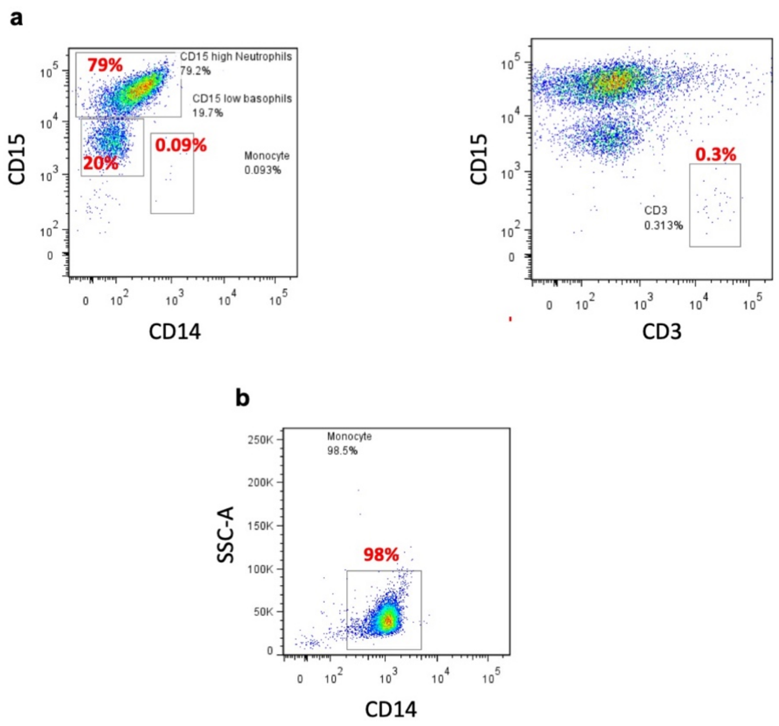


**Extended Data Fig. 8. Flow cytometric validation of purity of sorted cells from PBMCs. (a)** Purity of granulocyte cell populations. (**b**) Purity of monocyte cell populations. Antibodies: anti-CD15-FITC, anti-CD14-APC, and anti-CD3-PECy7.

#### List of Supplementary Tables

##### Supplementary Table 1. (separate file)

List of ChIP-seq samples, DA peaks, RNA-seq samples, DE genes, SEs and the GC-corrected and input-corrected peak heights for each ChIP-seq sample.

##### Supplementary Table 2. (separate file)

Gene Ontology analysis: functional enrichment of genes near DA peaks and enrichment of DE genes.

##### Supplementary Table 3.

Transcriptomic data sets used for *KCNJ15* differential expression analysis.

| ID | Country of cohort | Healthy controls | Active TB | Post-treatment active TB | Study reference | Microarray/ RNA-seq platform | GEO accession | Sample type |
|---|---|---|---|---|---|---|---|---|
| **UK '10 test set** | United Kingdom | 12 | 21 | - | Berry et al., 2010 | Illumina Human HT-12 v3.0 | GSE19444 | Whole blood |
| **UK '10 training set** | United Kingdom | 12 | 13 | - | Berry et al., 2010 | Illumina Human HT-12 v3.0 | GSE19439 | Whole blood |
| **TG '11** | The Gambia | 37 | 46 | - | Maertzdorf et al., 2011 | Illumina Human HT-12 v4.0 | GSE28623 | Whole blood |
| **UK '10 treatment set** | United Kingdom | - | 12 | 7 | Berry et al., 2010 | Illumina Human HT-12 v4.0 | GSE19435 | Whole blood |
| **Singapore monocytes** | Singapore | 19 | 20 | - | This study | RNA-seq Illumina HiSeq 2000 | This study | monocytes |
| **Singapore granulocytes** | Singapore | 18 | 20 | - | This study | RNA-seq Illumina HiSeq 2000 | This study | granulocytes |

**Supplementary Table 4. (separate file)**

The granulocyte and monocyte haQTLs.

**Supplementary Table 5. (separate file)**

The eQTLs and haQTLs from previous publications in LD with haQTLs from this study. The list includes all SNPs without collapsing the eQTLs by LD as done in the enrichment analysis in Figs. 5c,d and Extended Data Fig. 7.

**Supplementary Table 6. (separate file)**

The GWAS SNPs in LD with haQTLs. The list includes all SNPs without collapsing the haQTLs by LD as done in the enrichment analysis in Fig. 5e,f.

**Supplementary Video 1. (separate file)**

**Localization of Kir4.2 to BCG-containing lysosomes**. mcherry-BCG infected THP-1 monocytes were stained for lysosomes and KCNJ15/Kir4.2 24hr post-infection. Images were acquired using 3D-SIM. Scale bars: 2μm.
